# The representational geometry of emotional states in basolateral amygdala

**DOI:** 10.1101/2023.09.23.558668

**Authors:** Pia-Kelsey O’Neill, Lorenzo Posani, Jozsef Meszaros, Phebe Warren, Carl E. Schoonover, Andrew J.P. Fink, Stefano Fusi, C. Daniel Salzman

## Abstract

Sensory stimuli associated with aversive outcomes cause multiple behavioral responses related to an animal’s evolving emotional state, but neural mechanisms underlying these processes remain unclear. Here aversive stimuli were presented to mice, eliciting two responses reflecting fear and flight to safety: tremble and ingress into a virtual burrow. Inactivation of basolateral amygdala (BLA) eliminated differential responses to aversive and neutral stimuli without eliminating responses themselves, suggesting BLA signals valence, not motor commands. However, two-photon imaging revealed that neurons typically exhibited mixed selectivity for stimulus identity, valence, tremble and/or ingress. Despite heterogeneous selectivity, BLA representational geometry was lower-dimensional when encoding valence, tremble and safety, enabling generalization of emotions across conditions. Further, tremble and valence coding directions were orthogonal, allowing linear readouts to specialize. Thus BLA representational geometry confers two computational properties that identify specialized neural circuits encoding variables describing emotional states: generalization across conditions, and readouts lacking interference from other readouts.

## INTRODUCTION

In everyday life, we commonly ascribe emotional states to individuals by observing their responses to sensory stimuli or events. These responses include various behaviors like approach or avoidance, as well as physiological changes such as autonomic reactivity. In fact, sensory stimuli that elicit emotional states can cause individuals to generate not just one but multiple different responses, like freezing and fleeing to safety in response to a threat^1,2^. Although the sensory cues that elicit these responses often predict rewarding or aversive outcomes, similar responses can be observed in novel situations, highlighting the propensity of emotional behavior to generalize^3^. Neural mechanisms that underlie emotional behavior must therefore provide the capacity to assign emotional significance to sensory stimuli, to account for emotional generalization, and to connect representations of stimuli to circuitry that implements multiple behavioral and physiological responses used to ascribe emotional states to individuals. The underlying neural circuit mechanisms that satisfy these requirements of emotional processing remain unclear.

Prior studies have established the importance of the amygdala in mediating many aspects of emotion by combining lesions or silencing of neural activity with behavioral assays of emotional states^4-6^. More recent work has sought to identify the neural circuits within the amygdala that implement specific emotion-related operations by characterizing the response selectivity, projection target(s) and/or molecular transcriptomics of individual neurons^7-12^. Together these studies have led to the proposal that the basolateral amygdala (BLA) possesses distinct neural circuits specialized to encode the appetitive and aversive valence of stimuli, with each circuit projecting to different targets involved in generating particular valence-specific responses^7,9,11.13-19^. Empirical support for this proposal has been provided by experiments employing projection-specific, activity-dependent, and/or genetic strategies to perturb activity in targeted amygdalar neural circuit elements during assays of emotional behavior ^11,20,21^.

Although these studies have provided significant insight into valence processing, neuroscientists still face at least two formidable challenges for understanding the role of the amygdala in emotion. First, emotional experience and behavior are not fully explained by valence, as emotional states are far more rich, complex and nuanced than can be described by valence alone^3^. In short, if neural circuits in the amygdala only encode valence, variables describing many other aspects of emotional states would not be represented. Second, increasing data reveal that BLA neurons often modulate their activity in relation to two or more variables, a property known as ‘mixed selectivity’^22-24^ . In non-human primates, in addition to the valence of a conditioned stimulus, the activity of neurons in the amygdala can be modulated by stimulus identity and location, by the relative size of expected rewards, by un-cued and thereby abstract contexts, and by unconditioned stimuli of both valences in an expectation-dependent manner^25-28^. Mixed selectivity has also been described in studies of neural activity in rodent amygdala, though it has not been explored as extensively^29-30^. The complexity of amygdalar response properties poses a conundrum for the ability to generalize a readout. A decoder trained to read out valence in one context might not necessarily work in another context, as the neural representation of valence could be affected by all the other variables that define a context. Given that valence alone cannot describe all aspects of an emotional experience, individual neurons with complex response properties might actually participate in two or more neural circuits that represent distinct variables describing different aspects of emotional states.

Here, we employ a virtual burrow assay^31^ in head-fixed mice to investigate how the amygdala represents variables used to ascribe emotional states to individuals. We presented aversive and neutral conditioned stimuli to mice while either silencing or recording BLA neural activity. Aversive stimuli, more frequently than neutral stimuli, elicit two responses from mice in the burrow: tremble, which is akin to freezing, and ingress into the burrow enclosure, which is akin to a flight response. Tremble and ingress thereby correspond to behavioral responses that can be used to ascribe feelings of fear and safety to mice^1,2,32,33^. We observed that BLA inactivation eliminated differential responses to aversive and neutral stimuli, but did not eliminate the responses themselves. These data suggest that BLA transmits valence signals, not motor commands. However, individual neurons in BLA rarely represented only stimulus valence. Instead, their activity was also modulated by stimulus identity, and/or by the behavioral state of the mouse, as defined by tremble and ingress responses. Due to the heterogeneous and seemingly disorganized response properties of BLA neurons, we turned to analyses of representational geometry to illuminate the computational properties of recorded ensembles, and to thereby identify specialized neural circuits in terms of these properties. This approach overcomes the ambiguity of interpreting the activity of individual neurons when they exhibit mixed selectivity, and the consequent difficulty in assigning individual neurons to particular circuits based on their selectivity alone.

Analyses of representational geometry examine the configuration of points in the ‘activity space’. In this space, ‘points’ correspond to the activity of every recorded neuron in an ensemble plotted against each other for every experimental condition^27,34^. The configuration constitutes the representational geometry of a neural ensemble. In our analyses, we trained linear decoders to read out task-relevant variables from ensemble activity as defined by experimental parameters and observable behavioral measures. Training the decoder assigns a weight to each neuron for each variable being decoded, and determines the threshold (the hyperplane within the geometry) that best discriminates between the different conditions (values) of a variable. The assigned weights can be thought of as corresponding to the synaptic weights of recorded neurons onto target neurons that compute an output^35,36^. The vector orthogonal to the hyperplane defines a ‘coding direction’ for the readout of that variable. Neurons possessing non-zero decoding weights together determine the coding direction.

In this paper, we have used analyses of BLA representational geometry to answer three questions concerning the role of the the amygdala in representing emotional states. First, how does the amygdala represent more than just valence? Second, can amygdala ensemble activity account for emotional generalization? And third, can analyses of representational geometry be used to identify specialized neural circuits for encoding variables describing emotional states? Our results reveal that recorded BLA ensemble activity - despite the prevalence of mixed selectivity neurons - can realize particular representational geometries conferring two computational properties that together enable identification of specialized neural circuits for reading out variables describing emotional states. One property – realized when two variables are represented with orthogonal coding directions which results in a lack of interference between the two readouts – explains how one underlying BLA neural ensemble can represent more than just valence. The second property – realized when a variable is represented with lower-dimensional structure so that readouts of the encoded variable do not depend upon the values of other variables – explains how BLA ensemble activity can support emotional generalization across different conditions.

## RESULTS

### Aversively conditioned stimuli cause mice to tremble and ingress into the burrow

Mice underwent a Pavlovian aversive conditioning paradigm using four distinct olfactory conditioned stimuli, two that predicted footshock (CS+s), and two other stimuli that were not reinforced (CS-s) (Fig. 1A,B). Following a conditioning session, the four odors were presented randomly to head-fixed mice in a virtual burrow assay; neither the CS- or CS+ was paired with reinforcement during these experiments. Linear displacement of the burrow - its forward and backward movement - was measured with a laser sensor. During most of the testing session, mice displace the burrow negligibly (less than 0.5mm). We refer to such epochs as ‘stationary’. However, interspersed with stationary epochs, there are epochs in which mice induce small amplitude displacements that fluctuate with regular periodicity. We refer to these epochs as ‘tremble’. In other epochs, mice induce a large amplitude forward displacement that draws the burrow over them, which we refer to as ‘ingress’ (Fig. 1C). While tremble mirrors a freezing-like state (see below), ingress is a flight-like behavior into the protected environment of the burrow^31^.

**Figure 1.**
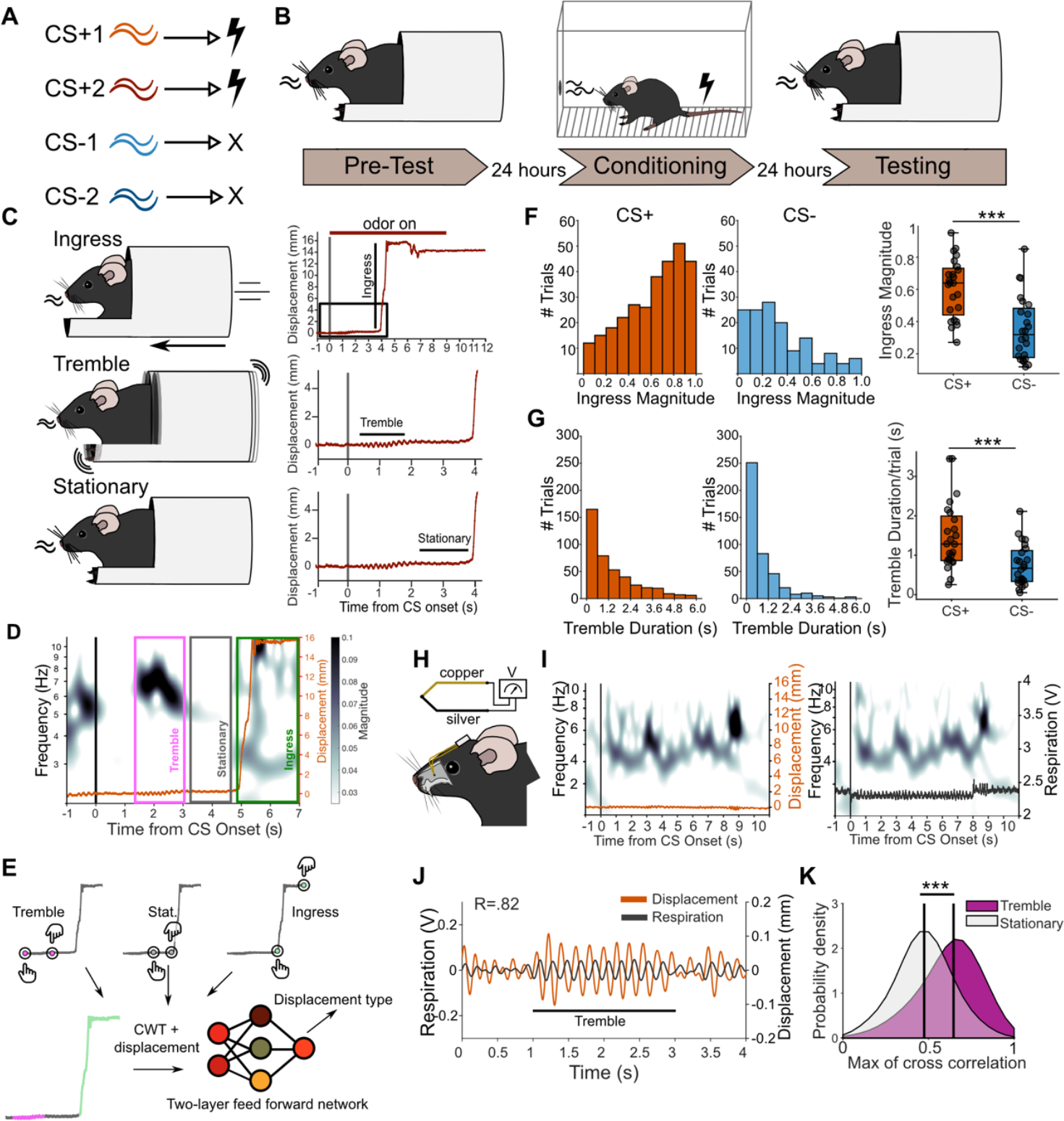
Aversive conditioned stimuli elicit tremble and ingress into a virtual burrow. **A**, Mice undergo Pavlovian aversive conditioning which employed four initially neutral odors. During conditioning, two odors are paired with footshock (CS+1 and CS+2), and two odors are unpaired (CS-1 and CS-2). **B**, Time course of experiment. Left, Pre-Test Session: all four odors are delivered unpaired to mice in the virtual burrow. Middle, Conditioning Session: 24 hours after Pre-Test, CS+ and CS-odors are presented in random order, CS+ odors co-terminate with footshock. Right, Testing Session: 24 hours after conditioning, the four odors are again presented to mice in the virtual burrow. **C**, Mice display three behaviors in response to odor presentation in the Testing Session. Top left, A large forward displacement of the burrow enclosure constitutes ‘Ingress’. Top right, burrow displacement as a function of time relative to CS+ onset from one trial in one mouse. Rectangle captures the time window depicted in zoomed-in graphs below. Middle left, A low amplitude oscillation of burrow position constitutes ‘Tremble’. Middle right, zoomed in view of displacement as a function of time with tremble period labeled. Bottom left, An epoch in which burrow displacement is negligible constitutes ‘Stationary’. Bottom right, zoomed in view of displacement as a function of time with stationary period labeled. **D**, Example cwt scalogram (frequency vs. time) showing 4-8Hz tremble, stationary and ingress periods in one CS+ trial. **E**, Diagram of neural network trained to predict tremble, stationary or ingress periods in the displacement data from a subset of labeled trials. **F**, Normalized ingress magnitude for each trial in response to CS+ (left) and CS-(middle) odors. Boxplots (right) show median normalized ingress magnitude in CS+/-for each mouse (n=24 mice, ***p<0.001, paired t-test). **G**, Same as (F) for tremble duration. Boxplot (right) is median tremble duration in CS+/-for each mouse (n=24 mice, ***p<0.001, paired t-test). **H**, Schematic showing thermocouple implant in nasal cavity to measure respiration in mice. **I**, Example scalograms of displacement (orange, left) and respiration (black, right) from one trial in one mouse. **J**, Example displacement and respiration during transition from a stationary to a tremble period (R=0.82 between tremble and respiratory oscillation during labeled ‘Tremble’ period). **K**, Probability density distributions of the maximum correlation between displacement and respiration during tremble (magenta) and stationary (light gray) periods (vertical lines depict median of distributions, tremble vs. respiration: 0.63, stationary vs. respiration: 0.4, ***p<0.001, Kolmogorov-Smirnov test).

We developed an unbiased approach for classifying the displacement data by using a wavelet transform to decompose the data into frequency components over time. Tremble periods are enriched in spectral power within the 4-8 Hz range (Fig. 1D). By comparison, an ingress, which appears as a rapid, step-like shift in displacement, predictably contains spectral power within all frequencies (Fig. 1D). To automatically label each frame as either tremble, ingress, or stationary displacement across experiments, we trained a neural network to classify every timepoint as one of the three categories (Fig. 1E). This classification allowed us to calculate, with millisecond resolution, the magnitude and the onset and offset times of both tremble and ingress responses, substantially extending previous characterization of mouse behavior in the virtual burrow^31^.

The presentation of CS+ odors increased the probability that a mouse would ingress (average number of ingresses: CS+, 12/20 trials; CS-, 6/20 trials; n=24 mice, p<0.001; Suppl. Fig. 1A). We computed the magnitude of ingress as the maximum displacement into the burrow attained on a trial relative to the maximal possible ingress (where a magnitude of 1.0 is maximal ingress). Across mice, ingress magnitudes on CS+ trials were higher than on CS-trials (CS+: 0.61; CS-: 0.35; n=24 mice, p<0.001; Fig. 1F). Furthermore, ingress on CS-trials was characterized by more fluctuations in displacement after ingress, and a logistic function therefore fit ingress less well on CS-than CS+ trials (CS+ r2: 0.85, CS-r2: 0.74, n=24 mice, p<0.01; Suppl. Fig. 1B). Meanwhile, tremble duration, maximum frequency, and average number of trembles per trial were all increased in response to a CS+ relative to a CS-(average time trembling per trial CS+: 1.48s, CS-: 0.75s; max frequency: CS+: 5.4Hz, CS-: 5.0Hz; average number of trembles: CS+: 0.93, CS-: 0.61, n=24 mice, p<0.001; Fig. 1G, Suppl. Fig. 1D,E). The latency to ingress or tremble did not significantly differ between CS+ and CS-trials (ingress onset CS+: 3.5s CS-: 3.1s, tremble onset CS+: 2.3s CS-: 2.4s; n=24 mice, p > 0.05; Suppl. Fig. 1C,F). Additionally, there was no significant difference between the spectral power of tremble on CS+ and CS-trials (CS+: 3.3, CS-: 2.7; n=24 mice, p> 0.05; Suppl. Fig. 1G).

In behavioral paradigms in which mice are not head-fixed, freezing is accompanied by the modulation of autonomic responses such as respiration and heart rate^37–41^. To assess whether trembling, like freezing, is related to changes in breathing, we implanted a thermocouple in the mouse nasal cavity to monitor changes in respiration (Fig. 1H). Breathing rate during tremble was within the 4-8Hz frequency range, overlapping the frequency range of displacement changes in tremble (Fig. 1I,J Suppl. Fig. 2A,B). Spectrograms of breathing and tremble oscillations were remarkably phase-coupled, and tremble and respiration bouts were highly correlated (tremble vs. respiration: r=0.63, stationary vs. respiration: r=0.4, p<0.001; Fig. 1K, Suppl. Fig. 2C). The tight correlation between breathing and tremble suggests that the displacement fluctuations during tremble may reflect inertial motions due to inhalation-exhalation cycles of sufficient strength to appreciably shift the animal’s center of mass. The minimal friction of the burrow assay enables us to capture these displacements. Overall, these data are consistent with studies in freely moving mice that have demonstrated a relationship between freezing elicited by an aversively conditioned stimulus and respiration^38,39^. Trembling therefore bears similarity to freezing, which is normally thought of as the absence of movement, and not a coordinated motor action. In a previous study, optogenetic activation of BLA neurons responsive to footshock were shown to modulate breathing and heart rate, a finding consistent with autonomic reactivity caused by changes in a mouse’s emotional state^8^. Indeed, activating the same footshock-responsive neurons can induce associative fear learning^8^.

**Figure 2.**
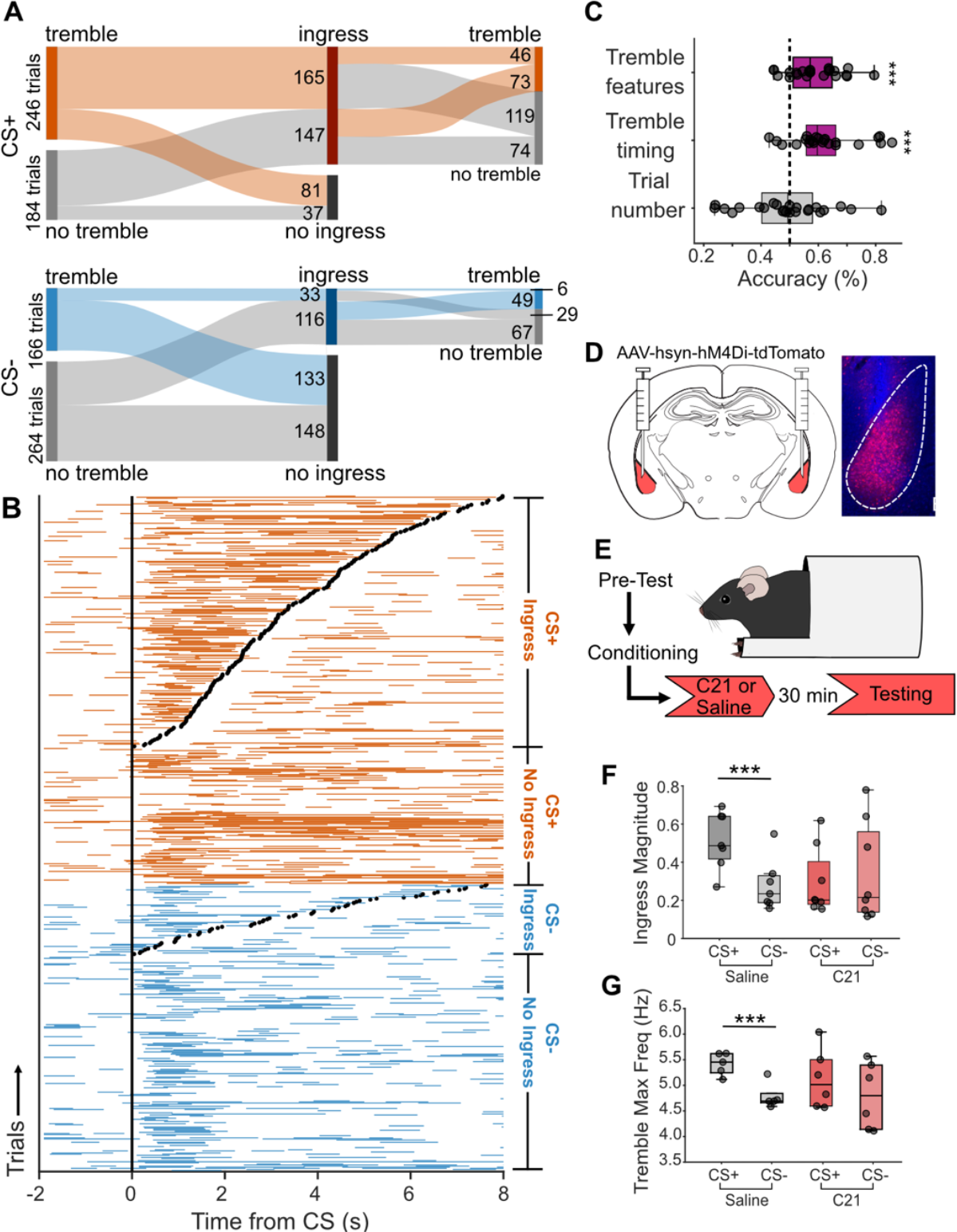
BLA activity is required for differential expression of tremble and ingress. **A**, Alluvial diagram showing the relationship between tremble and ingress for CS+/- (860 total trials, 430 trials per CS type, n=24 mice). **B**, Tremble bout(s) on ingress and no ingress trials for CS+ (orange) and CS- (blue) odors. Trials with ingress are ordered based on ingress onset time (dots) following CS onset (vertical line). **C**, Logistic regression models trained to classify trembles that transition to ingress within 1 sec of tremble offset vs. other trembles not immediately followed by ingress. Models include either tremble features (maximum frequency, spectral power), tremble timing variables (duration, number of trembles on trial, tremble latency), or trial number. Box plots depict median accuracies across mice (tremble features: 58%, tremble timing: 64%, n=24 mice, dashed vertical line indicates 50% chance accuracy, *** p<.001, t-test from chance accuracy). **D**, Bilateral injections in BLA of an adeno-associated virus harboring the inhibitory DREADD hM4Di and the fluorophore tdTomato (AAV-hsyn-hM4Di-tdTomato). Example injection site in one mouse, white dashed line outlines BLA. **E**, Prior to testing, mice were given an i.p. injection of the DREADD agonist, Compound 21 (C21), or saline (n=8 mice C21, n=7 mice saline). **F**, Box plots showing normalized ingress magnitude (CS+/saline: 0.51, CS-/saline: 0.28; CS+/C21: 0.29, CS-/C21: 0.34) and tremble maximum frequency (CS+/saline: 5.32Hz, CS-/saline: 4.73Hz; CS+/C21: 5.14Hz, CS-/C21: 4.82Hz) to CS+ and CS-in Saline and C21 groups (***p<.001, paired t-test).

According to predator imminence theory, an animal’s defensive strategy shifts from freezing to flight behaviors as the animal’s perception of threat escalates^42-44^. Analogously, although tremble and ingress possess distinct dynamics, trembles largely occur prior to ingress on individual CS+ trials. When a mouse trembled on a CS+ trial, ingress followed tremble on 51% of trials (165/319 tremble CS+ trials). In contrast, only 15% of trembles on CS-trials were followed by ingress (33/215 tremble CS-trials; Fig. 2A,B). Note that odors are presented for 8 seconds, so both tremble and ingress invariably occur during the constant delivery of a conditioned odor (Fig. 2B). Consequently, mice transition between tremble and ingress responses without changes in external cues.

Given the observation that ingress tends to follow tremble, we wondered whether measurable aspects of a tremble could predict a subsequent ingress. We trained a decoder to distinguish between trembles that transitioned to ingress within 1 second or less of tremble offset compared to trembles that did not lead to ingress. Features related to tremble quality (the maximum frequency of the tremble and spectral power) or to the timing of trembles (tremble latency, duration, and the number of trembles in a trial) accurately predicted transition to ingress on a trial at above chance levels (58% accuracy using tremble features, p<0.001; 64% accuracy using timing, p<0.001, n=24 mice; Fig. 2C). Trial number was not predictive of ingress (51% accuracy, p>0.5, n=24; Fig. 2C). Since ingress can be predicted by specific features of a tremble bout, tremble and ingress are linked behaviors, analogous to freezing followed by flight to a safe environment. The virtual burrow assay therefore enables us to measure these linked behaviors that can be used to ascribe emotional states without using compound stimuli or explicit training^18,45,46^.

### BLA inactivation eliminates differential tremble and ingress responses to aversive and neutral stimuli

We hypothesized that readouts of activity in BLA neural ensembles would be required for mice to exhibit differential tremble and ingress responses to CS+ and CS-presentations. We therefore tested whether decreasing BLA activity alters these differential responses by using a pharmacogenetic approach to inhibit BLA activity during presentations of odors in the burrow. An AAV (adeno-associated virus) harboring the inhibitory DREADD (Designer Receptor Exclusively Activated by Designer Drug) receptor hM4Di and tdTomato was bilaterally injected into BLA (Fig. 2D). Prior to testing in the burrow, we separated mice into two groups and administered the DREADD agonist Compound 21 (C21) or saline through intraperitoneal injection (Fig. 2E). In animals injected with C21, ingress magnitude was diminished in response to the CS+, becoming similar to levels observed in response to the CS-(ingress magnitude CS+/saline: 0.51, CS-/saline: 0.28, n=7, p<0.001; ingress magnitude CS+/C21: 0.29, CS-/C21: 0.34, n=8, p>0.05; Fig. 2F). Similarly, the maximum frequency of tremble during the CS+ was significantly reduced in C21 mice (max frequency CS+/saline: 5.32Hz, CS-/saline: 4.73Hz, n=7, p<0.001; max frequency CS+/C21: 5.14Hz, CS-/C21: 4.82Hz, n=8, p>0.05; Figure 2f). Thus, like freezing and avoidance responses in freely moving animals^47,48^, both differential tremble and ingress responses to a CS+ and CS-depend on activity in the BLA. Despite these results, it is unlikely that BLA provides a motor signal that generates tremble and ingress responses. Trembling and ingress still occur following BLA inactivation; it is differential responses to the CS+ and CS-that are eliminated. These results are consistent with studies showing that amygdala neurons do not exhibit activity time-locked to responses elicited by appetitive or aversive conditioned stimuli^13^.

### BLA ensemble activity encodes the tremble and ingress response states

Tremble and ingress responses define two different states that can be used to ascribe fear and safety to mice. We next sought to determine whether BLA neural ensembles provide a representation of these distinct states. We used two-photon calcium imaging to measure cellular activity in the BLA while odors were presented to a subset of mice (Fig. 3A, Suppl. Fig. 3). Neural activity from states defined by tremble or whether mice had ingressed into the safety of the burrow were each compared to activity during states when mice were stationary in the burrow and not in the ingressed position. All of these states occur within the 8 sec odor delivery. Note that since mice are head-fixed in the virtual burrow, the proximity of the nose to the odor port is the same across states.

**Figure 3.**
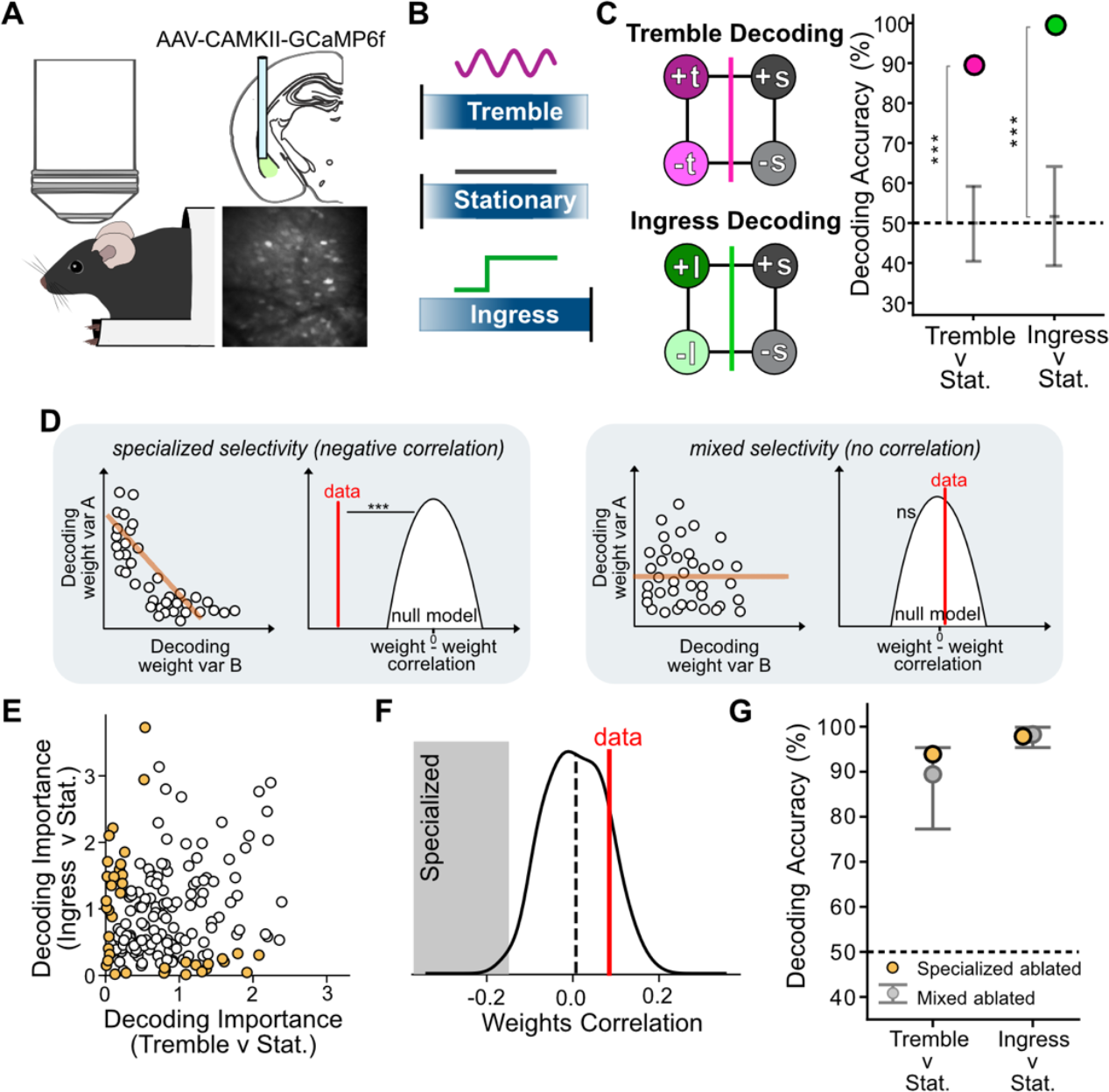
Mixed selectivity in BLA: encoding of tremble and ingress response states. **A**, Schematic of two-photon imaging setup during testing in the virtual burrow assay (left), schematic depicting location of viral injection site and GRIN lens in BLA (top right), and imaging field of view in one mouse (bottom right). **B**, Neural activity during tremble, stationary and ingress epochs were selected for decoding analyses. Tremble and ingress epochs were chosen from different trials. **C**, Left: a linear decoder is trained to decode the tremble (top) or ingress (bottom) states compared to the stationary state, with stimulus valence balanced across conditions (+t or +i or +s, tremble, ingress or stationary epochs from CS+ trials; -t or -i, or-s, tremble, ingress or stationary epochs from CS-trials). Right: decoding accuracy for tremble or ingress vs. stationary conditons. Grey error bars, 95% confidence intervals of the null model (see Methods). *** p < 0.001. **D**, Illustrations depicting how the Spearman correlation coefficient can be used to determine if a neural ensemble exhibits more specialized selectivity than expected by chance. Left, scatterplot of toy data where the absolute value of decoding weights for two variables is plotted against each other. In this example, neurons tend to have high decoding weights for one or the other variable, but not both (specialized selectivity). The regression slope (orange line) and Spearman correlation coefficient are negative. The observed Spearman correlation coefficient is compared to the distribution of correlation coefficients obtained from a null model that implements mixed selectivity in the neural ensemble (see Methods). If there is more specialized selectivity than expected by chance, the observed correlation coefficient (red line, ‘data’) will be less than 95% of the values obtained from the null model. Right, scatterplot of different toy data where many neurons exhibit high decoding weights for both variables. There is no correlation between decoding weights for the two variables (orange line), and the Spearman correlation coefficient (red line, ‘data’) falls within the null distribution, indicating that in this case there is not more specialized selectivity than expected by chance (mixed selectivity). **E**, For each recorded neuron, the absolute value of its decoding weights for tremble and ingress are plotted against each other. Yellow data points, neurons with apparent ‘specialized’ selectivity that are artificially ablated in (**G**). **F**, Spearman’s correlation from comparison of decoding weight data depicted in (**E**) (red line, ‘data’) and null model (distribution) (null: 0.01±0.07, data: 0.08, p=0.13). **G**, Decoding performance for tremble and ingress (vs stationary) after ablating neurons with specialized selectivity (**E**, yellow data points), or with mixed selectivity (averaged across iterations). Grey error bars, 95% confidence interval for null model generated from iterations of ablating different, random selections of mixed selectivity neurons.

We used a linear classifier to determine if the tremble and ingressed states are each decodable from ensemble activity as compared to the stationary state. Analyses focused on neural activity during three time epochs: trembling bouts that lasted at least 500ms (“tremble”), the 2-second time epoch after mice had ingressed into the burrow (“ingress”), and time epochs of at least 500ms in which mice were stationary and had not ingressed into the burrow (“stationary”) (Fig. 3B). Since tremble and ingress occur more commonly on CS+ compared to CS-trials, we balanced the conditions included for decoding by subsampling the data such that decoding performance could not derive from correlations between CS valence and tremble or ingress. The conditions used for decoding tremble or ingress response states therefore contained a 50-50 blend of CS+ and CS-trials for the trembling, ingressed and stationary conditions (Fig. 3C).

Both being in the tremble and ingress states compared to a stationary state were robustly decodable across the neuronal population and across individual mice (tremble 90% accuracy; ingress 100% accuracy; p<0.001; Fig. 3C, Suppl. Fig. 4A). Decoding performance was asymptotic when ∼50% of neurons were included (Suppl. Figure 4B). The ability to decode whether mice are in the tremble, ingress, or stationary state did not depend on movements involving different magnitudes of displacements of the burrow across the three states. When we selected tremble and ingress bouts where the displacement range of the burrow was minimal (less than 0.5mm, Suppl. Fig 4C), both states remained highly decodable (tremble decoding: 90% accuracy, p<0.001; ingress decoding: 94% accuracy, p<0.001; Suppl. Fig. 4D). In addition, the ability to decode tremble was not due to the fact that trembles occur more frequently prior to an ingress. If we considered only trials where mice later ingressed, decoding accuracy for tremble vs. stationary states was 100% (p<0.001; Suppl. Fig. 4E). Thus, even though ingress and tremble are correlated, and even though a tremble often leads to an ingress, a neural representation of the tremble state exists independent of the ingress state.

**Figure 4.**
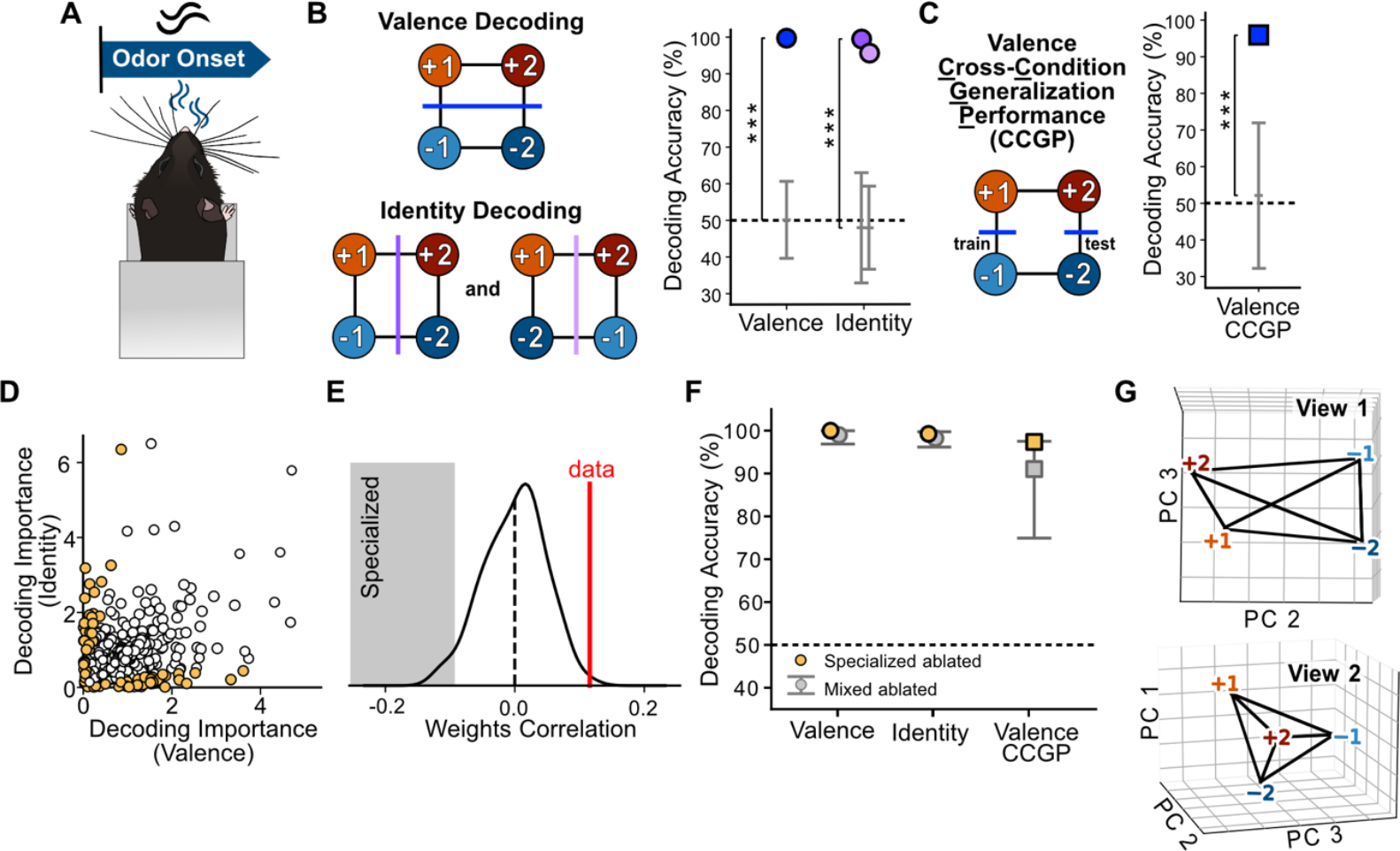
BLA representational geometry encoding CS valence and identity following CS onset. **A**, Neural activity during the 2-second window of time following CS onset was selected for decoding analysis. **B**, The four CS conditions (+1, + 2, -1, and -1, corresponding to CS+1, CS+2, CS-1, CS-2) were divided into two equally sized groups in three ways (three dichotomies). The CS valence variable describes one dichotomy; the other two dichotomies relate to the encoding of CS identity. A linear classifier is trained to distinguish between the conditions on each side of a dichotomy using randomly selected trials from all conditions. The classifier is then tested on held-out trials. Decoding accuracy for stimulus valence, and for each dichotomy related to CS identity was high (valence: 100% accuracy, identity dichotomies: 99% and 96%; ***p<0.001; error bars, 95% confidence interval of null model). **C**, Cross Condition Generalization Performance (CCGP) involves training a decoder on trials from only a subset of conditions, and testing on trials from held-out conditions, as exemplified schematically for the variable valence (e.g. +1 v. -1 for training; +2 v. -2 for testing, see Methods). CCGP for valence was high (96% accuracy, ***p<0.001; error bars, 95% confidence interval of null model). **D**, For each neuron, the absolute value of its decoding weight for valence is plotted against its decoding weight for one of the two identity dichotomies shown in (**B)**. Yellow data points, neurons with apparent ‘specialized’ selectivity neurons that were artificially ablated in (**F). E**, Spearman’s correlation calculated between the decoding weights for valence and identity (red line, ‘data’) and null model (distribution) (null: -0.00±0.05, data: 0.12, **p<0.01). **F**, Decoding performance for valence, and one of the identity dichotomies as well CCGP for valence after ablating neurons in (**D)** with apparent specialized selectivity (yellow), or mixed selectivity (averaged across iterations of ablating randomly selected mixed selectivity neurons). Grey error bars, 95% confidence interval for null model generated from iterations of ablating mixed selectivity neurons. **G**, Two views of PCA plot depicting BLA representational geometry following CS onset. Top, visualization of parallel coding directions across pairs of conditions for CS valence, which accounts for high CCGP. Bottom, visualization showing that despite possessing low-dimensional structure for encoding valence, the geometry has higher dimensional components that enables decoding of CS identity.

In principle, decoding the tremble and ingress states from a stationary state could be based on the activity of distinct BLA sub-populations, one specialized for the tremble state and the other for the ingress state. In this case, neurons would exhibit specialized selectivity for one or the other response state, but not both. To assess the contribution of neurons with mixed as opposed to specialized selectivity for readouts of the tremble and ingress states, we used two approaches to quantify the selectivity of each recorded neuron. One approach used the population decoding weight of a neuron as a proxy for its selectivity for the variable being decoded, and the other approach used a non-parametric method, ROC analysis, to characterize selectivity based on firing rates inferred from calcium signals of single neurons^13^. ROC analysis computes the area under the ROC curve (AUC), with larger AUC values indicating more selectivity. Of course, trembles and ingress did not occur at the same time, so this analysis examined data from the same neurons at different timepoints.

Once we had quantified the selectivity of each recorded neuron, we computed three different statistical measures that were then used to quantify the degree to which specialized selectivity was observed across the population. The three measures were applied to both metrics of selectivity (ROC analysis and decoding weights) and involved calculating 1) the Spearman correlation coefficient between selectivity measures for both variables; 2) a mean specialization index that quantified the relative selectivity between the two variables; and 3) an index called the L-index for quantifying the degree to which datapoints within a 2-dimensional scatterplot of selectivity for the 2 variables lie along the axes (see Methods). For all three statistical measures, we determined if the observed value was indicative of more specialized selectivity than expected given a null model that implements mixed selectivity across the population. Briefly, the null model is created by performing iterations of a solid random rotation of observed ensemble activity in the neural activity space, breaking the alignment of neural responses with specific neural axes and thus implementing mixed selectivity. For example, a highly negative Spearman correlation coefficient between selectivity measures for two variables would indicate that neurons that strongly encode one variable are typically not relevant for the other variable (Fig. 3D, left panel). In this case, if an observed Spearman correlation coefficient is less than 95% of the values of the null model, we conclude there is more specialized selectivity than expected by chance. By contrast, when no correlation is observed between selectivity measures, the observed data would fall within the distribution of correlation coefficients from the null model, indicative of a predominance of mixed selectivity neurons (Fig. 3D, right panel). A positive correlation between selectivity measures would also indicate a predominance of mixed selectivity in the ensemble.

When we applied these methods to the data used to decode tremble and ingress, the scatterplots of decoding weights and AUCs for both variables revealed that many neurons exhibited a high degree of activity in relation to both response states (Fig. 3E; Suppl. Fig. 4F). The Spearman correlation coefficient, selectivity index and L-index values all indicated that neurons did not exhibit more specialized selectivity than expected by chance, whether using decoding weights or ROC analyses to quantify selectivity (weights correlation, Fig. 3F; weights selectivity, weights L-index, Suppl. Figure 4E; AUC correlation, selectivity and L-index, Suppl. Fig 4F). Moreover, artificial ablation of the neurons that appeared to exhibit specialized selectivity for variables describing the tremble or ingress state (i.e. eliminating from analyses neurons that were close to either axis in the selectivity plot, Fig. 3E) had minimal effects on decoding performance, indistinguishable from artificial ablations of a similar number of mixed selectivity neurons (p > 0.05, Fig. 3G).

Given that the same neural ensemble – comprised of neurons exhibiting mixed selectivity could be read out to decode both the tremble and ingress states, we wondered whether the brain states corresponding to these two states were distinct. We found that we could decode the tremble compared to the ingress states from each other using BLA ensemble activity (81% accuracy, p<0.01). Consistent with this finding, a linear classifier trained to decode tremble vs. stationary states did not generalize to decode ingress vs. stationary states (52% accuracy, p>0.05). This finding indicates that the coding direction for distinguishing between the tremble and stationary states is not parallel to the coding direction distinguishing between ingress and stationary states, as parallel coding directions would support cross-state generalization of the trained decoder^27,34^.

### BLA representational geometry following CS onset: mixed selectivity and the encoding of valence

The fact that mice tremble and ingress more frequently to CS+ as compared to CS-stimuli suggests that a computation of the valence of stimuli impacts the propensity to generate tremble and ingress responses. To determine if BLA neural ensembles encoded the valence of odor stimuli in the burrow, we analyzed two-photon calcium imaging data within a 2 second window following CS onset (Fig. 4A). Our experimental design used four different odor CSs (CS+1, CS+2, CS-1, and CS-2; Fig. 1A). A variable describing valence corresponds to a dichotomy that divides these four conditions into two equally sized groups (CS+1/CS+2 vs. CS-1/CS-2, Fig. 4B). Training and testing a classifier on this dichotomy resulted in high decoding accuracy for valence both across the BLA neuronal population, and on average across individual mice (population: 100% accuracy, p<0.001; average individual subject’s decoding accuracy: 74%, p<0.05; Figure 4B, Suppl. Fig. 5A). The encoding of valence was robust, as decoding performance became asymptotic once only ∼25% of randomly selected neurons were included in the analysis (Suppl. Fig. 5B). However, valence is not the only information represented in BLA during this time. The four CS conditions can be divided into two other equally sized dichotomies (CS+1/CS-1 vs. CS+2/CS-2, and CS+1/CS-2 vs. CS+2/CS-1; Fig. 4B). Variables describing each of these other dichotomies could also be decoded at high levels (99% and 96% accuracy, p<0.001; average individual subject’s decoding accuracy: 65%, p<0.05; Fig. 4B, Suppl. Fig. 5A). This analysis indicates that BLA ensemble activity differs for all four CS conditions. As a result, CS identity is represented in BLA, consistent with observations made previously in studies in monkeys^13,24,49^.

**Figure 5.**
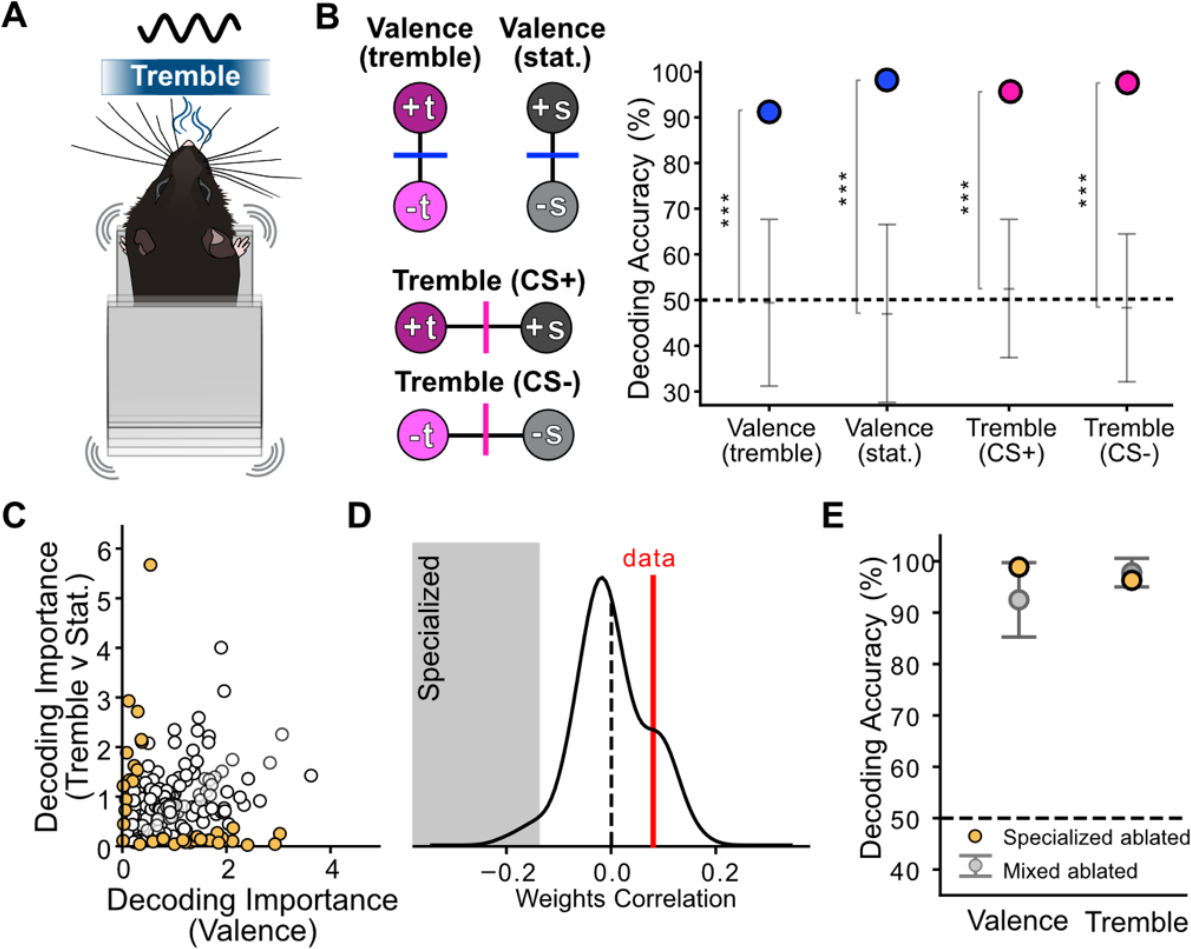
Mixed selectivity for tremble and valence in BLA. **A**, Neural activity within time epochs of at least 500 ms during tremble or stationary states was selected for decoding analyses. **B**, Trial conditions were defined by response state and stimulus valence (+t, +s: tremble, stationary on CS+ trials; -t, -s: tremble, stationary on CS-trials). Decoding performance was high for valence on both tremble and stationary trials and decoding performance was high for tremble on both CS+ and CS-trials (valence (tremble): 92% accuracy, valence (stationary): 98%), tremble (CS+): 96%, tremble (CS-): 98%; ***p<0.001; grey error bars, 95% confidence intervals for the null model). **C**, For each neuron, the absolute value of its decoding weight for valence is plotted against its decoding weight for tremble when decoding each variable across all four conditions. Yellow data points, neurons with apparent ‘specialized’ selectivity that are artificially ablated in (**E**). **D**, Spearman’s correlation coefficient calculated from the data in (**C**) (red line, ‘data’) and null model (distribution) (null: 0.00±0.07, data: 0.08, p=.12). **E**, Decoding performance for valence and tremble vs. stationary after artificially ablating neurons with apparent specialized selectivity (yellow) in (**C**), or with mixed selectivity (average performance across ablation iterations). Grey error bars, 95% confidence interval for null model generated from iterations of ablating mixed selectivity neurons.

Novel stimuli, and not just previously experienced ones, can elicit approach or defensive responses, even when the novel stimuli are neutral. This phenomenon is called generalization, a cardinal feature of emotions observed across species^3^. To our knowledge, no prior work has established whether the representational geometry of stimulus valence within BLA can in principle support this type of generalization. We used a recently described analytic approach, cross-condition generalization performance (CCGP)^27^, to determine if the observed representational geometry in BLA could support the generalization of valence across different stimuli. To compute CCGP for valence, a decoder is trained to classify CS+ vs. CS-using only 1 exemplar of each type of CS condition (e.g. CS+1, CS-1), and then tested on held out types of conditions (e.g. CS+2, CS-2) (Fig. 4C). CCGP is then defined as the average decoding performance across all ways of choosing the testing and training CS conditions^27^.

High CCGP is observed only with certain representational geometries, in particular, geometries in which pairs of conditions that correspond to two values of the same variable have parallel coding directions^27^. For the variable valence in our experiments, this type of geometry would be observed if the coding direction for pairs of conditions with opposing valences (CS+ vs. CS-) tend to be parallel. Indeed, in our data, during the 2-second interval after CS onset, CCGP for valence was significantly greater than chance as computed from the null model (96% accuracy; p<0.001 Fig. 4C). Since CCGP corresponds to the decoding accuracy on CS+ and CS-trial types that are not used for training, the data indicate that a decoder can classify a stimulus as a CS+ or CS-even if the stimulus is “new” to the decoder. This computational property indicates that the geometry of a neural representation of valence reflects a process of abstraction, a process that creates lower dimensional structure within the representational geometry^27^.

Conceivably, BLA neurons that encode CS valence could be specialized, representing valence equivalently for all stimuli. This type of specialized selectivity for a variable could account for high CCGP^27^. We therefore assessed the selectivity of recorded neurons across the population for CS valence and CS identity. Scatterplots of decoding weights and AUC values for the valence and identity variables revealed a prevalence of neurons that exhibited some degree of selectivity for both variables (Fig. 4D, Suppl. Fig. 5D). The observed Spearman correlation coefficients, mean specialization indices, and L-indices all indicated that the population was comprised predominantly of mixed selectivity neurons, with specialized selectivity not being observed more often than expected by chance (Fig. 4E, Suppl. Fig. 5C,D). Results were similar regardless of which identity dichotomy was employed. Moreover, decoding performance and CCGP for valence did not depend on neurons with specialized selectivity. Artificial ablation of those neurons that appeared to exhibit specialized selectivity for variables describing valence and one of the identity dichotomies, which entails removing those neurons from decoding analyses, had minimal effects on decoding performance or CCGP, and the effects were statistically indistinguishable from artificial ablations of a similar number of mixed selectivity neurons (p > 0.05; Fig. 4F).

Together these results indicate that the representational geometry in BLA following CS onset has two main features. The geometry has a lower-dimensional structure with respect to encoding valence, accounting for high CCGP. At the same time, the representation allows all stimulus identities to be decodable. This representational geometry can be visualized using a dimensionality reduction technique. The visualization enables one to appreciate that the coding directions for CS valence (i.e. the lines connecting different CS+ and CS-conditions) tend to be parallel so as to confer high CCGP for valence across stimuli (View 1, Fig. 4G). However, the overall geometry is not planar and all binary variables describing the four CS conditions can be read out (View 2, Fig. 4G). This higher dimensionality allows for input-output computations specific for the identity of stimuli.

### BLA ensembles comprised of mixed selectivity neurons encode tremble and valence

The analysis of tremble behavior revealed that it occurs more commonly after an aversive as compared to a neutral odor presentation, indicating that tremble responses are based in part on an evaluation of the valence of a stimulus. We therefore sought to determine if variables describing aversive vs. neutral stimulus valence and tremble vs. stationary response states were represented by BLA ensembles. We again analyzed neural activity when mice were in the tremble and stationary states by selecting time epochs lasting at least 500 ms for each (Fig. 5A), with an equal number of CS+ and CS-trials included for analysis in both tremble and stationary states. Decoding of CS valence was robust in both the tremble and stationary states considered individually, as was decoding of tremble when considering CS+ and CS-trials individually (valence decoding accuracy = 92% and 98% for tremble and stationary states; tremble decoding accuracy = 96% and 98% for CS+ and CS-trials; p<0.001 in all, Fig. 5B). These results indicate that BLA neural ensembles do not only represent stimulus valence, and nor do ensembles only represent tremble vs. stationary states.

Whether characterizing neuronal selectivity using decoding weights or AUC methods (Fig. 5C, Suppl. Fig. 6B), all measures of specialization (Spearman correlation coefficient, mean specialization index, and the L-index) indicate that across the recorded neural ensemble there was not more specialized selectivity for valence and tremble than expected by chance (Fig. 5D, Suppl. Fig. 6A,B). Nonetheless, valence as well as tremble were robustly decodable across the neuronal population (valence 95% accuracy, tremble 98% accuracy, p<0.001). Excluding from analysis those neurons whose decoding weights suggested specialization for either valence or tremble did not diminish decoding performance, and the effects of such artificial ablations did not differ significantly from ablations of an equivalent number of neurons with mixed selectivity (p>0.05; Fig. 5E).

**Figure 6.**
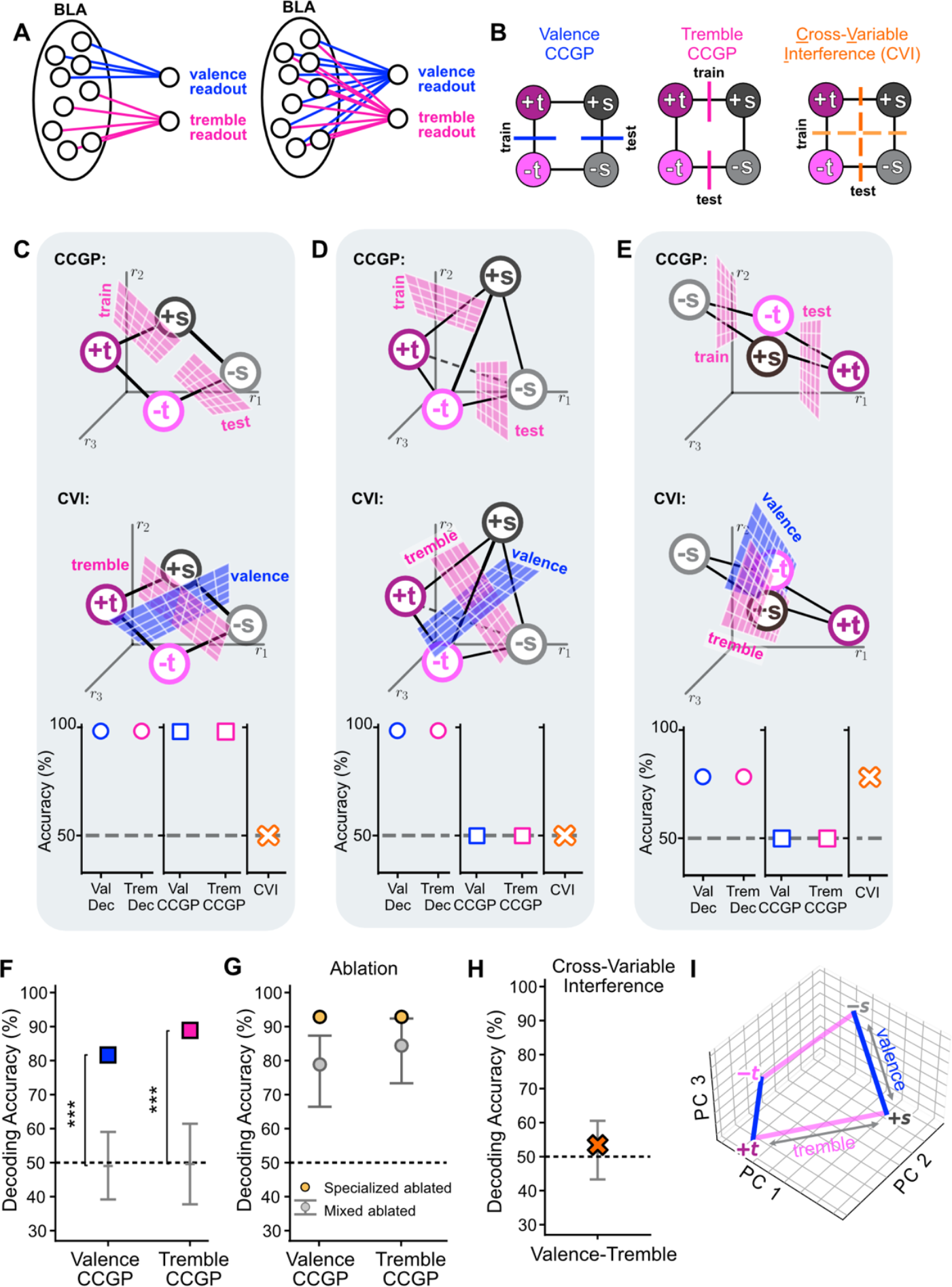
BLA representational geometry reveals specialized and disentangled circuits for tremble and valence. **A**, Left, simple neural network comprised of neurons with specialized selectivity for valence or tremble. Each neuron contributes to the readout of either valence or tremble, but not both. Right, neural network comprised of neurons with mixed selectivity for valence and tremble. Each neuron contributes to the readout of both valence and tremble. **B**, Two measures of representational geometry used to characterize the computational properties of the neural ensemble: CCGP (computed for valence and tremble), and cross-variable Interference. To compute cross-variable interference, a linear classifier is trained to decode a variable (e.g. valence) using randomly selected trials from all 4 trial conditions (tremble and stationary on CS+ and CS-trials). During training, each epoch of activity is labeled according to valence. The decoder is then tested on held-out trial conditions where each epoch of neural activity is labeled according to response state, ‘tremble’ or ‘stationary’ and not according to valence. When decoding performance is at chance, the coding directions for the two variables are orthogonal (no interference between coding directions). When CCGP is high and cross-variable interference absent, neural circuits for each readout are specialized. **C-E**, Different representational geometries confer different computational properties to neural circuits encoding valence and tremble. Plots depict geometries when the activity of three mixed selectivity neurons is plotted in the firing rate space across four trial conditions. **C**, A geometry where tremble and valence are decodable, CCGP is high for both variables (top, CCGP analysis shown for tremble), and there is no interference between readouts of the two variables (bottom). Here the coding directions for tremble and valence are orthogonal with respect to each other (see hyperplanes for decoding the two variables). **D**, A representational geometry where the response at each condition is at a random location in the firing rate space (forming a tetrahedron). Tremble and valence are decodable, but generalization across conditions is at chance (top). There is no cross variable interference (bottom). Thus in some geometries, coding directions can be orthogonal, but generalization across conditions is not conferred. Circuits encoding variables in this geometry are not specialized. **E**, A representational geometry where generalization across conditions is not conferred (top), and there cross-variable interference between readouts is present (bottom). In this case, cross variable interference is due to the fact that the coding directions for tremble and valence are parallel. **F**, BLA neural ensembles exhibit a geometry where CCGP for valence and tremble is high (valence CCGP: 82% accuracy, tremble CCGP: 89%; ***p<0.001; grey error bars, 95% confidence intervals for the null model). **G**, Decoding accuracy after artificial ablation of neurons with apparent specialized selectivity or after artificial ablations of equivalent numbers of mixed selectivity neurons (grey dot, average; grey error bars, 95% confidence interval for null model generated from iterations of ablating mixed selectivity neurons). **H**, Decoding performance for cross-variable interference is at chance (accuracy: 54%, p>0.05). **I**, BLA representational geometry visualized with dimensionality reduction. The coding directions for tremble (pink lines) and valence (blue lines) appear to be orthogonal (explaining the absence of cross-variable interference), yet the coding directions for pairs of conditions for each variable appears parallel, accounting for high CCGP.

### Distinct specialized circuits for tremble and valence

We next sought to determine if specialized neural circuits encode aversive vs. neutral stimulus valence as compared to the tremble vs. stationary response states. In principle, simple specialized circuits for reading out valence and tremble could be comprised of neurons in BLA that are specialized (Fig. 6A, left). In this case, one subpopulation of neurons could encode only valence, and another subpopulation could encode only tremble. A valence readout can be readily constructed by connecting one or more output neurons only to those BLA neurons encoding valence. Analogously, neurons specialized to encode tremble would comprise a distinct specialized circuit. These types of simple circuits have at least two advantageous computational properties: 1) the two readouts do not interfere with each other, as the tremble state does not affect readouts of valence, and tremble readouts do not depend on valence; and 2) each readout circuit can easily generalize to novel situations, as the activity of output neurons trivially depends only on one variable. However, these kinds of simple specialized circuits for representing valence and tremble are not compatible with what we observed in BLA neural ensembles, as neurons predominantly exhibited mixed selectivity for these variables.

Even if neurons predominantly exhibit mixed selectivity, there are representational geometries that allow one to construct specialized circuits for readouts of valence and tremble (Fig. 6A, right). These circuits can possess the same computational properties as the simple circuits just described. To quantify these properties, we use two measures of representational geometry, CCGP and a new measure called cross-variable interference (Fig. 6B, see Methods). When CCGP is high, a readout circuit can generalize to novel situations, and when the value of one variable does not affect the readout of another variable, there is no cross-variable interference between readouts of these two variables. Together these properties can define specialized neural circuits even when neurons exhibit mixed selectivity.

To illustrate how analyses of representational geometry can reveal specialized neural circuits, we consider three different geometries that each encode the tremble and valence variables (Fig. 6C-E). In the depictions of these geometries, the activity of three mixed selectivity neurons are plotted against eachother in the firing rate space for four different conditions defined by whether mice were in the tremble or stationary state on either CS+ or CS-trials. One geometry confers high generalization performance across conditions for both variables (high CCGP for valence and for tremble), and has no cross-variable interference (Fig. 6C). The lack of cross-variable interference indicates that the coding directions for tremble and valence are orthogonal, as a decoder trained to readout one variable is at chance when tested on decoding the other variable. Since generalization across conditions is high and cross-variable interference is absent, the neural circuits for representing both tremble and valence possess computational properties that define specialized circuits. The contribution of any given neuron to a specialized circuit is determined by its decoding weight. In a different geometry, where the activity for each condition is plotted in a random location in the firing rate space, both tremble and ingress can be robustly decoded with little cross-variable interference. However, generalization across conditions for both variables is at chance (Fig. 6D). Thus the circuits are not specialized. Finally, in geometries where the coding directions for tremble and valence are nearly parallel, cross-variable interference is high, and generalization across conditions is low (Fig. 6E). In this case, cross-variable interference is high because the coding directions for tremble and valence are parallel. When this occurs, the same hyperplane that separates tremble from stationary response states also can classify valence at above chance levels.

Given this conceptual framework for understanding how representational geometry confers computational properties, we analyzed BLA ensemble activity to determine if tremble and valence are represented in specialized circuits. First we discovered that the representational geometry of tremble supports cross condition generalization across valence, and the representional geometry of valence generalizes across tremble and stationary response states (CCGP for valence: 82%; CCGP for tremble: 89%; p < 0.001 in both cases, Fig. 6F). These generalization properties did not depend upon neurons with apparent specialized selectivity, as artificial ablation of such neurons did not diminish CCGP and did not differ from artificial ablation of equivalent numbers of mixed selectivity neurons (Fig. 6G). The results indicate that the representational geometry has low dimensional structure in representing valence across tremble vs. response states, and in representing response state across stimulus valence. Moreover, during the tremble state, the representation of valence also has low dimensional structure with respect to CS identity, as CCGP for valence across stimulus identity is high during the tremble state, just like during the epoch following CS onset (86%, p<0.001). These findings were observed despite the fact that mixed selectivity for CS identity and valence was prevalent, with CS identity being decodable at above chance levels (71%, p<0.01).

Given that tremble and valence were represented with a geometry that supported generalization across conditions, we next determined if the readouts for tremble and valence exhibited little cross-variable interference. We observed an absence of interference, as a decoder trained to decode valence is at chance for decoding tremble (and vice-versa for a decoder trained to read out tremble)(decoding accuracy = 54%, p > 0.05, Fig. 6H). The coding directions for tremble and valence are therefore orthogonal (no interference). Fig. 6I provides a visualization of the recorded representational geometry using dimensionality reduction. The coding directions for valence appear to be parallel across response states, and the coding directions for tremble vs. stationary response states appear to be parallel across both valences of stimuli. These geometric properties account for the observed high cross condition generalization for both variables. At the same time, coding directions for valence and tremble appear to be approximately orthogonal in a quite planar geometry (similar to the schematic in Fig. 6C). Thus despite the prevalence of mixed selectivity neurons - whereby the same neurons often have non-zero decoding weights for readouts of both variables - the observed BLA geometry confers two computational properties that define specialized circuits for tremble and valence: generalization across conditions and an absence of cross-variable interference.

### BLA representational geometry reveals a circuit for safety

Ingress, like tremble, occurs more frequently after a CS+ than a CS-presentation, and BLA inactivation eliminates this differential behavior (Suppl. Fig 1A,E; Fig. 2A,E,F). These results indicate that an ingress response is based in part on an assessment of stimulus valence. However, stimulus valence may become behaviorally irrelevant once mice have ingressed into the safety of the burrow. We hypothesized that the representation of stimulus valence in BLA in fact would depend upon whether mice had ingressed into the safety of the burrow. Neural activity during the 2 seconds before and the 2 seconds after an ingress (Fig. 7A) were analyzed. As before, we subsampled trials so that an equal number of CS+ and CS-trials were included for all analyses. We discovered that stimulus valence was decodable immediately before an ingress, but not right afterwards (accuracy prior to ingress = 96%, p < 0.001; accuracy after ingress = 41%, p > 0.05; Fig. 7B). BLA therefore encoded stimulus valence in a manner dependent on the response state. The representation of valence was not inextricably elicited by the presentation of a conditioned stimulus.

**Figure 7.**
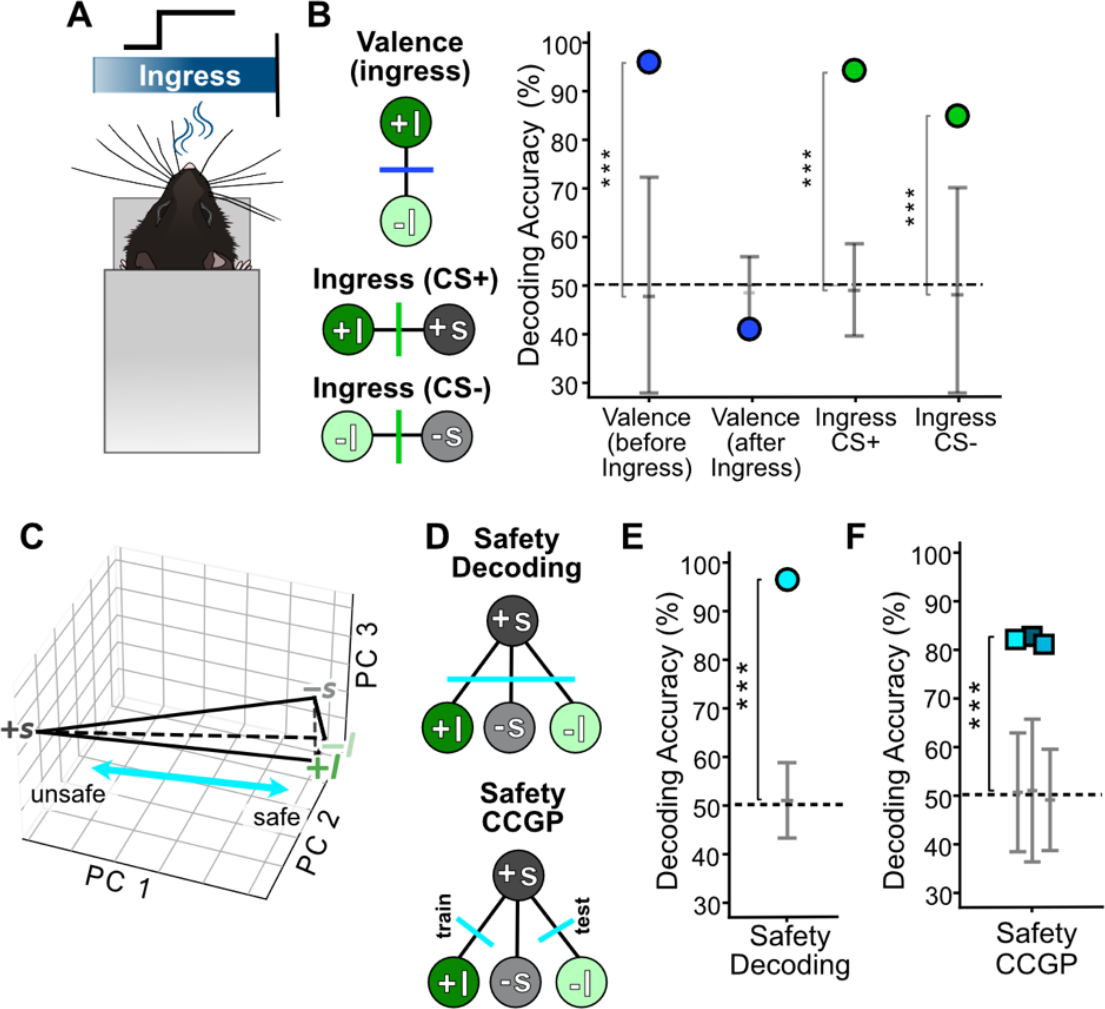
BLA representational geometry of safety. **A**, Neural activity is analyzed in three time epochs: a 2-second time window after mice reach the ingressed position, a 2-second before they ingress, or when they are stationary and have not ingressed. Since odor delivery remains constant during these time periods, conditions used for decoding can be defined by whether the animal will or has ingressed (I) and whether the odor present is a CS+ or CS-. **B**, Valence is highly decodable before ingress but not right after ingress (accuracy prior to ingress = 96%, p < 0.001; accuracy after ingress = 41%, p > 0.05). Meanwhile, ingress is decodable on CS+ and CS-trials (Ingress CS+: 94%, Ingress CS-: 85%, ***p<0.001). **C**, PCA plot depicting BLA representational geometry during ingress and stationary conditions (trial conditions labeled according to ingress/stationary and valence: I, ingress; s, stationary; +, CS+; -, CS-. **D**, Dichotomies for examining the encoding of safety, and for evaluating if the encoding of safety generalizes across different types of conditions. For safety, the variable being decoded is the +S condition vs. the three remaining conditions (top). CCGP for safety is calculated by adapting our standard CCGP approach. A decoder is trained to decode safety using half the +S (unsafe) trials from one of the safe trial conditions (+I, -I, or -S; bottom diagram illustrates training on +S vs. +I). The decoder is then tested on decoding safety using the remaining half of +S trials from each of the remaining two safe trial conditions (bottom panel depicts testing on held out +S trials vs. -I trials). This CCGP analysis is performed 3 ways so that each of the safe trial conditions is used for training. **E**, BLA neural ensemble activity encodes safety (safety decoding: 96%, ***p<0.001). **F**, CCGP for safety is high (safety CCGP: 82%, 83%, 81%, respectively, for each of the 3 ways of training the decoder; ***p< 0.001).

Although valence is not decodable immediately after an ingress, BLA neural ensemble activity did encode whether mice had ingressed into the burrow, or not, on both CS+ and CS-trials (Fig. 7B). Within BLA representational geometry, the coding directions for ingress and valence were not orthogonal, as there was some cross-variable interference (Suppl. Fig. 7A). This does not imply that the representation of ingress and valence in BLA is identical, as it is not. For example, the encoding of valence does not generalize across ingress and stationary states, but encoding of ingress does generalize across stimuli of both valences (Suppl. Fig. 7B).

We sought to gain more insight into BLA representational geometry in the ingressed vs. stationary conditions by using dimensionality reduction. This visualization suggested that the pattern of activity on CS-trials when mice were stationary and not ingressed is similar to the activity observed when mice are in the ingressed state (regardless of CS valence). This pattern of activity is quite different from that observed when a CS+ is presented and mice are stationary (not having ingressed) (Fig. 7C). This observation raises the possibility that a variable other than ingress or valence might describe low dimensional structure in BLA representational geometry during this time interval.

Within our experiment, three conditions exist that might allow one to conclude that a mouse feels “safe”: one in which a neutral stimulus (CS-) is presented and mice have not chosen to ingress and therefore remain stationary, and two other conditions, CS+ and CS-trials when mice have chosen to ingress inside the burrow. By contrast, an unsafe condition could be experienced by a mouse upon presentation of a shock-associated CS+, prior to their executing an ingress. Thus a variable we refer to as safety divides trials into two (uneven) groups: +S (unsafe) trials, and +S/+I/-I (safe) trials (Fig. 7C).

To determine whether and how BLA neural ensembles represent safety, we quantified the computational properties of BLA representational geometry using decoding and an adaptation of our CCGP analysis (Figs. 7D). These analyses revealed that BLA ensemble activity could be used to decode with high accuracy the +S (unsafe) condition vs. the other three safe conditions combined (96%, p<0.001; Fig. 7E). Furthermore, the variable safety was generalizable across different pairs of conditions (mean accuracy across the three ways of creating pairs of safe/unsafe conditions: 82%, p < 0.001; Figure 7F). In these analyses, a linear classifier that separates half of the +S trials from the trials on one of the safe conditions successfully separates the remaining +S trials from the trials of each of the remaining safe conditions. Indeed, the coding directions between the +S (unsafe) condition and each of the three safe conditions tended to be parallel, unlike the coding directions for separating any of the other conditions from the remaining 3 conditions (Suppl. Fig. 7C). Safety was therefore represented by BLA ensembles with low dimensional structure, just as depicted Fig. 7C. This representational geometry can account for the emotional generalization of safety across different types of conditions.

## DISCUSSION

Emotional states are commonly ascribed to individuals by observing their responses to sensory stimuli and events. In this paper, we presented aversive and neutral stimuli to mice in a virtual burrow assay, eliciting two responses used to infer distinct states: tremble for fear, and ingress into the burrow for safety. Inactivation of BLA did not eliminate tremble and ingress, but did eliminate differential responses to aversive compared to neutral stimuli. These results are consistent with the notion that BLA signals stimulus valence, not motor commands. Two-photon imaging data of cellular activity was then used to understand how the BLA encodes stimulus valence, as well as response variables used to ascribe emotional states to mice.

Across the population of BLA neurons recorded, neurons predominantly exhibited mixed selectivity for multiple variables, not specialized selectivity for only one variable like valence. The seemingly disorganized and heterogenous pattern of response selectivity, whereby the diversity of coding properties is vast, suggests that specialized circuits in BLA may not be identified by only considering single neuron properties such as physiological response selectivity, projection patterns, and/or molecular transcriptomics. Instead, we propose that analyses of representational geometry – analyses that take into account neural activity across a neural ensemble and across experimental conditions - provide a new approach for identifying specialized circuits in BLA by characterizing the computational capabilities conferred by recorded geometries.

Our analyses of representational geometry reveal that BLA ensemble activity possesses distinct coding directions for three variables describing elements of emotional state: valence, a fear state defined by the tremble response, and a variable describing whether mice are in a safe state, which occurs when they ingress into the burrow or when a CS-appears and they choose not to ingress. BLA representational geometry was highly structured with respect to how it encoded these three variables, despite the fact that neurons predominantly exhibited mixed selectivity. Linear readouts of valence, tremble and safety did not depend on the value of other variables, indicative of lower-dimensional structure in BLA representational geometry for encoding these variables. This lower-dimensional structure enables generalization across different conditions, a computational property that could underlie different forms of emotional generalization^3^. Finally, stimulus valence and the tremble response state are simultaneously represented by BLA, with orthogonal coding directions for the two variables, which means a low level of cross-variable interference between linear readouts of the variables. The lack of interference combined with the capability of generalizing across conditions indicate that valence and tremble are each encoded by specialized BLA neural circuits. However, the same individual neurons often participate in both specialized circuits, as evidenced by the fact that neurons predominantly exhibit mixed selectivity and have non-zero decoding weights for readouts of both variables. Indeed, for all of these computational properties, individual neurons exhibiting apparent specialized selectivity for any one of the variables were not required. Eliminating such neurons from analyses did not diminish decoding performance or the ability to generalize across conditions.

### On valence and functional circuits in BLA

In prior studies, individual neurons in BLA have often been observed to exhibit selective responses to the valence of either or both conditioned and unconditioned stimuli^8,11,13,19,50^. These results have suggested the existence of specialized circuits for valence^51^. In support of this, a number of studies have demonstrated that perturbing the activity of specific BLA neurons or their nerve terminals at projection targets can cause valence-specific effects on emotional behavior^10,11^. Further, appetitive and aversive neural circuits in BLA were recently described as being comprised of anatomically segregated and genetically defined populations of BLA neurons: Rspo2-expressing pyramidal neurons and another population expressing Ppp1r1b^9^. However, subsequently, it has been shown that Rspo2-expressing neurons are not homogenous, and subpopulations of these neurons have different valence preferences depending upon their projection target^12^. This raises the possibility that a combination of response selectivity, anatomical projection target and molecular transcriptomic characterization can identify the neurons that participate in valence-specific BLA neural circuits. Most studies, however, have not characterized neural responses properties across a broad enough range of parameters to assess the degree to which neurons exhibit mixed selectivity. Despite the relatively small number of conditions in our experiments, by measuring neural activity across conditions and across the evolving responses to stimuli, our data demonstrate that mixed selectivity is prevalent in BLA. This observation is consistent with increasing numbers of studies that have emphasized the complexity of physiological response properties of BLA neurons^24,30,52,53^. Mixed selectivity neurons can contribute to computations of multiple different outputs, indicating that such neurons could participate in multiple specialized circuits, not just one.

If an individual BLA neuron participates in multiple specialized circuits, then this could be mediated by different anatomical pathways. BLA neurons have been shown to project to multiple anatomical targets^10,54^, and these neurons can have different synaptic weights at each target. Since inactivation of BLA results in the elimination of differential tremble and ingress responses to aversive and neutral odors without eliminating the responses themselves, BLA neurons may transmit valence information to different brain targets that implement tremble and ingress. Relevant to our experiments, genetically defined, distinct circuit elements have been identified in CeA that mediate freezing and fleeing to safety in freely moving animals^46^. These responses are analogous to the tremble and ingress responses observed when odors are presented to mice in the burrow assay. Physiological and optogenetic evidence has shown that activity in somatostatin-expressing (SOM+) neurons is related to the generation of freezing, while activity in corticotropin-release-factor–expressing (CRF+) neurons mediates flight behaviors. While BLA input to SOM+ and CRF+ neurons in CeA has been demonstrated^46,55^, it remains unknown if individual BLA neurons project to both of these distinct neural populations. However, regardless of whether such direct projections exist, polysynaptic pathways may exist both within and beyond the amygdala. For example, BLA neurons project to the intercalated masses, to the insula, and to numerous other structures that then have direct connections to CeA^56,57^. In addition, complex processing within BLA may occur within its microcircuitry, as recent experiments show that valence-selective BLA neurons inhibit some nearby neurons and excite others^54^. These locally activated and inactivated neurons may have various response properties and projection targets, providing another means for individual neurons to contribute to more than one output computation. Advances in multiphoton techniques that allow the simultaneous perturbation and recording of activity-defined ensembles in BLA may help shed light on mechanisms at the microcircuit level that explain how mixed-selectivity neurons can be part of multiple specialized circuits^50^.

The fact that the inactivation of BLA eliminates differential tremble and ingress to aversive compared to neutral stimuli suggests that output neurons for implementing each tremble and ingress likely receive input about valence from BLA either directly or, as just discussed, indirectly. However, the implementation of a response to a stimulus with valence can also modulate the representation of valence itself. We observed that valence encoding in BLA is state-dependent, disappearing upon ingress despite the continued presence of conditioned odors. Similar state-dependent modulation of valence representations have also been observed in non-human primate amygdala^26^. Thus, additional mechanisms (and associated anatomical pathways) must exist that actually alter the representation of valence in BLA. These mechanisms are likely different from mechanisms mediating extinction. Conditioned odors do not lose their potency on trials that follow trials in which mice have ingressed into the burrow. On trials subsequent to an ingress, a CS+ still elicits ingress at higher rates than a CS-.

Although understanding how valence representations in BLA can be modified is important for understanding the role of the amygdala in emotional behavior, our data indicate that valence coding alone does not fully account for how the amgydala represents emotional states. The amygdala also represents the tremble vs. stationary response state while representing valence. As we have shown, the pattern of breathing observed during tremble is similar to that observed when freely moving mice freeze. Freezing is traditionally used to ascribe fear to mice, and thus tremble may also be used to ascribe a fear state. Many responses that reflect an underlying emotional state, including tremble and freezing, actually fluctuate after the presentation of an emotionally significant stimulus. If the BLA had only encoded valence, and not also tremble, it would not have represented aspects of the richness and complexity of emotional states that can be revealed by examining activity in relation to responses that dynamically evolve.

Finally, and critically, in the BLA the coding direction for tremble vs. stationary states is approximately orthogonal compared to the coding direction for CS valence, as there is an absence of cross-variable interference in readouts. Moreover, the representations of tremble and valence each generalize across the conditions of the other variable. Lack of interference and generalization across conditions therefore reveal specialized neural circuits for tremble and valence. Using terminology from the machine learning community, the representations of these variables are therefore ‘disentangled’^58^.

### Representational geometry and the generalization of emotional states

The generalization of emotional states across situations, including to new ones, has long been appreciated as an important phenomenon implicated in psychopathology^59-63^, but prior studies have typically focused on the role of the amygdala at the cellular/molecular level in stimulus generalization, not generalization across many types of conditions^64,65^. The capacity to generalize across situations relies on identifying their shared features. The process of identifying shared features across instances - a process called ‘abstraction’ - can provide a more compact and typically dimensionally reduced representational geometry^66,67^. However, if representations discard all details of the environment, fewer variables are represented, which in turn limits the number of input-output operations that can be performed (limiting the number of responses that can be generated by reading out activity)^22,23,27^. In other words, as an example, responses specific to stimulus identity cannot be generated if all information about stimulus identity is discarded and only valence information saved.

In our data, BLA representational geometry displayed low dimensional structure when encoding three variables: 1) CS valence (aversive vs. neutral stimuli) across the tremble vs. stationary responses states, and across stimulus identity immediately after CS onset and during the tremble state; 2) tremble vs. stationary response states across stimuli of both valences; and 3) unsafe vs. different types of safe states (ingressed into the burrow for stimuli of both valences, and a state when mice remain stationary after a CS-presentation). As a result, the same output can be generated across different types of inputs for each of these three variables. However, despite this ability to generalize, other variables were also often represented. CS identity is encoded at the same time as valence. Similarly, ingress vs. stationary states can still be readout when safety is represented and the geometry has a low-dimensional structure. These types of compromises between representing one or more variables in a manner that can support generalization across conditions while still being able to perform additional input-output computations have been observed previously in monkeys^27^.

To provide an intuition as to why high CCGP can account for emotional generalization, consider the representational geometry encoding the variable valence both after CS onset and during the tremble state. When an ensemble of BLA neurons responds to a novel stimulus, the pattern of activity will be located somewhere along the coding direction for valence. Depending upon the location along the valence coding direction, the novel stimulus will be decoded as either aversive or not, even though 1) it has no prior emotional meaning, and 2) that pattern of activity has never been activated in the past. In principle, a variety of factors could dictate where a novel stimulus is represented with respect to the valence coding direction in BLA. These factors include an individual’s pre-existing affective state, the similarity of a stimulus to other previously experienced stimuli possessing emotional significance, and the environmental situation. Regardless of the factors that might modulate how a novel stimulus is represented in BLA, the disentangled structure for encoding valence means that readouts of valence for a novel stimulus could produce emotional states and their associated valence-specific behavioral and physiological responses. Similarly, given high CCGP for tremble across stimulus valence, depending upon where the response to a stimulus, even a neutral or novel stimulus, lies along the coding direction for tremble vs. stationary states determines whether the animal will tremble or not. Finally, given high CCGP for safety, different types of safe conditions can be readout in the same way to generate across conditions the same types of safety-related responses, such as feelings and physiological responses associated with safety.

### Emotional states and neural circuit specialization through the lens of representational geometries

Three central questions have long hovered over studies of the neuroscience of emotion. How does the brain represent variables describing emotional states? How do such neural representations account for emotional generalization? And how does one identify specialized neural circuits for representing these variables? Our results indicate that analyses of the representational geometry of BLA neural ensembles provide a means of answering these questions. Neural circuit specialization is often conceptualized as relying on computations implemented by neurons specialized to encode information about only one variable or one particular combination of variables. Accordingly, in the amygdala, circuit specialization for representing variables describing emotional states has often focused on identifying neurons that encode information about valence[REFS]. However, here we observed that BLA neurons generally do not encode only valence, as they predominantly exhibit mixed selectivity for combinations of variables, including stimulus identity and response states defined by tremble and ingress in the virtual burrow. Nonetheless, BLA ensemble activity can realize a representational geometry that confers two vital computational properties commonly thought to rely on neurons encoding only a single variable. One property is achieved when the representational geometry possesses orthogonal coding directions for two (or more) variables. In this situation, there is no interference between readouts of the different variables, as was the case when considering the encoding of valence and tremble in our data. As a result, the same neural ensemble can simultaneously represent multiple variables describing emotional states in an independent manner, a richer representation than just encoding valence. The second property is achieved when a geometry encodes a variable with a low-dimensional structure, enabling generalization across different types of conditions (high CCGP), which we observed with respect to the encoding of valence, tremble, and safety. When both properties are satisfied, a variable is represented in a specialized circuit.

These findings form the basis of a conceptual framework whereby specialized circuits in BLA are not defined by the presence of neurons with selectivity for one variable or one combination of variables. Instead, neural circuit specialization is identified by considering activity across a neural ensemble and across conditions, using analyses of representational geometry to characterize computational capabilities. This conceptual framework is appealing because it no longer relies on the characterization of individual neurons alone to identify specialized neural circuits. Further, the framework explains how individual neurons that exhibit mixed selectivity and that may also project to more than one anatomical target can participate in multiple specialized circuits. The enhanced ability to record from large neural ensembles in the context of different behaviors, combined with analyses of representational geometry, thereby can provide a powerful tool for identifying specialized circuits that otherwise might have been elusive given the complexity of both neural response properties and anatomical connectivity that is commonly observed in many brain areas.

## Acknowledgements

We thank Richard Axel for his generosity and numerous contributions, as well as Joshua Gordon and members of the Salzman lab for helpful comments on the manuscript. PK.O. was supported by NIMH T32MH015144, BBRF Young Investigator Award, and NIMH K01MH123783. C.D.S. and S.F. were supported by R01-MH082017 and the Simons Foundation.

## METHODS

### Mice

Male C57BL/6J mice (Jackson Laboratories) were singly housed on a 12-h light-dark cycle with ad libitum access to food and water. All procedures were performed in accordance with standard ethical guidelines approved by Columbia University Institutional Animal Care and Use Committee. 24 mice were used for initial experiments characterizing behaviors. A subset of 9 mice underwent two photon imaging. A cohort of 15 mice were used for pharmacogenetic experiments. All experiments were performed during the light cycle on three consecutive days. Surgical procedures were performed on mice at 8-10 weeks of age. Mice included in all behavioral, inactivation and imaging experiments were between 12-16 weeks of age.

### Virus injection surgical procedure

Animals were anesthetized with isoflurane and secured in a stereotactic apparatus (Stoelting). Internal body temperature was regulated through a heating pad placed under the body. Ophthalmic ointment was applied to the eyes to prevent corneal drying (Patterson Veterinary) and Buprenoprhine (0.05mg/kg) was administered as an analgesic. The surgical site was clipped and cleaned with iodine followed by ethanol washes. Lidocaine was applied topically to the scalp to numb the area of incision. The skull was exposed and cleaned with sterile cotton swabs. One machine screw was fixed to the skull in order to anchor the dental cement. For behavior only experiments, the skull surface was then immediately covered in a thin layer of cement (MetaBond, Parkell Inc). A titanium head plate was lowered onto the skull, centered at lambda and secured with additional applications of cement.

For DREADD inactivation and calcium imaging experiments, following attachment of the machine screw, a craniotomy was performed using a microdrill to access the brain for viral injection. All injections were performed using a microinjection pipette, Nanoject II (Drummond 6563A01). For inactivation experiments, we made bilateral injections of 200-400 nl of hSyn-hM4Di-tdTomato (Addgene, 50475-AAV2) into the BLA (AP:+/-3.30, ML: -1.55, DV: -4.95, -4.85, relative to bregma). For calcium imaging experiments, we made a unilateral injection of 600nl CAMKII-GCaMP6f (Addgene, 100834-AAV1) in the left hemisphere using the same coordinates. Viral injections were followed either by headpost attachment or both headpost attachment and GRIN lens implantation. Mice were given 2 weeks to recover following headpost only surgery and 4 weeks to allow for adequate viral expression following viral injections before starting behavioral experiments.

### Implantation of GRIN lens

In the group of mice that underwent two-photon imaging, a 0.85mm microendoscope probe (Grintech GmBH) was stereotactically implanted in the BLA. Brain tissue was first aspirated to a depth of ∼1mm below skull surface to reduce intracranial pressure from the implant. The lens implant was lowered to DV -4.8mm relative to bregma and fixed in place using VetBond adhesive (3M) followed by a layer of dental cement and head post attachment. A layer of Kwik-cast (World Precision Instruments) was applied to the GRIN lens to protect it from damage between imaging sessions. After 4-6 weeks, the level of GcaMP6f expression was assessed using the two-photon microscope. Imaging experiments began when viral expression was sufficiently bright, as judged by visual inspection.

### Implantation of thermocouple

To measure respiration, one mouse was implanted with a custom-made thermocouple. Thermocouple wires were inserted in the intranasal cavity and affixed to the skull using dental cement.

### Virtual Burrow Assay

Detailed methodology of the virtual burrow assay design and operation is described in the original publication^31^. Briefly, the virtual burrow is composed of a 3D-printed tube enclosure (virtual burrow). A trimmed, absorbent underpad is attached to the floor of the burrow, underneath the mouse, and a wooden stick is adhered to the front of the tube to serve as a handlebar for mice to grip with their forepaws. The virtual burrow is mounted on near-frictionless air bushings set at 15 psi (New Way Air Bearings). These air bushings slide on two parallel rails to constrain movement in one dimension. The animal is head-fixed using two custom-machined clamps fitted to the titanium headplate cemented to the mouse skull. The burrow is tethered to a linear actuator with a fishing line. The actuator is employed to place the mouse in a neutral position within the burrow during intertrial periods. The linear displacement of the mouse is measured using a laser displacement sensor focused on a 3D-printed flag that is affixed to the side of the tube enclosure. The laser readout provides a continuous, time-dependent, one-dimensional variable of position. All analog voltage signals from the laser displacement sensor are digitized at 1kHz using a National Instruments NI-DAQ for behavior-only experiments, or at 10kHz for calcium imaging experiments using a Blackrock Microsystems DAQ.

### Odor delivery

Odorants acetophenone, octanal, pinene, and limonene (Sigma Aldrich) were mixed at a 1% dilution in mineral oil before the start of each experiment. Odorants were delivered through a custom-designed olfactometer. Custom-written Arduino code was used to structure the trials, which included communication between the virtual burrow assay and the olfactometer. Odor stimulus onsets were logged on the same DAQ as incoming burrow displacement data. Odor concentrations during each trial were recorded using a photoionization detector (PID) as voltage signals also collected on the DAQ.

### Aversive conditioning paradigm

During pre-test, mice in the virtual burrow assay were presented with odor stimuli in 10 blocks of trials. Each block consisted of the 4 odors delivered in a random order for a total of 40 trials. Odor stimulus duration was 8 seconds and odor identity was varied for different mice (e.g., acetophenone was a CS+ for one mouse and a CS-for another mouse). The next day, mice were placed in a Med Associates chamber fitted with a grid floor in order to deliver foot shocks. During conditioning, each odor was delivered 8 times, with two odors paired with a 0.7mA footshock while the other two remained unpaired. Odor delivery was controlled by an Arduino and footshock triggered with TTL pulses to Med Associates shock scrambler. The following day, mice were tested in the virtual burrow assay using the same block structure as pre-test day.

### Two-photon imaging

All images were acquired using a multi-photon microscopy system (Prairie Technologies, now Bruker) with a 20x objective (0.5 NA, Nikon) and an ultra-fast pulsed laser beam (920-nm wavelength, Coherent). Time series were collected at 6.5 Hz and 128x256 pixels per frame. We continuously acquired fluorescence with a photomultiplier tube (GaAsP PMT) operated by PrairieView software. Each mouse received two imaging sessions, during pretest and test days (all data presented is from test day).

### Extraction of calcium traces

Calcium imaging time-series were motion corrected using a planar hidden Markov model (SIMA)^68^. Data were then imported to MATLAB. Cell regions of interest (ROIs) were identified using the MATLAB CaImAn package^69^; spatial footprints and deconvolved signal for ROIs were extracted using CNMF.

### Neural network classification of behavior and respiration

We trained a neural network to distinguish between periods of stationary displacement, tremble displacement, and directed motion. The model was designed to perform the classification task of distinguishing between the three movement states based on displacement and spectrograms of the displacement. First, we randomly sampled subset of 100 trials, with equal contributions from each of the 24 mice and with both CS+ and CS-trials present in the subset. For these trials, we labeled segments of the displacement as either “stationary”, “tremble”, or “motion”. The labeled data was resampled to produce equal numbers of each of the three movement categories. These labels then served as the ground truth for training and evaluating the neural network model. As features for the model, we input the displacement, as well as the displacement convolved with kernels of varying sizes to capture past and future displacement, and lastly, values from each trial’s spectrogram computed over the displacement using MATLAB’s cwt (continuous wavelet transform) function. A two-layer feed-forward neural network was trained using MATLAB’s patternnet function. During training, the network learns to map the engineered features to the correct class labels. Categorical cross-entropy is used as the loss function. A subset of test data from a held-out set of trials is taken to validate the model. Model performance is evaluated using metrics including accuracy, precision, recall, and F1-score on the testing dataset. The trained neural network was then used to label every time point of every trial’s spectrogram and displacement data into “stationary”, “tremble”, or “motion”.

### Behavior and respiration quantification

The continuous wavelet transform was computed from the displacement and respiration data using MATLAB’s cwt function on a per-trial basis, with parameters set to 10 octaves and 48 voices per octave. Additionally, for each trial, “ingress onset” was defined as the first timepoint when displacement exceeded 1 mm. The “ingress magnitude” was calculated as a ratio of the maximum displacement attained following an ingress divided by the maximum achievable within the virtual burrow assay for the session. The “tremble duration” was computed as the total time from all the tremble segments in a trial.

### Prediction of ingress using tremble features

To explore a possible relationship between tremble and ingress, the trials were separated into ingress and no-ingress trials. Three logistic regression models were trained and tested using sklearn’s LogisticRegression module within Python with balancing of class weights. The three models were based on (1) tremble timing features including the duration of a segment, the time that segment began, and the total time spent trembling in that trial; (2) the tremble’s mean spectral power between 4-8 Hz and the tremble’s maximum frequency; and (3) the trial number on which a tremble occurred. The cross-validated accuracy was computed using 150 cross-validations for each model.

### Neural decoding analysis

Neural decoding analysis was performed using the Decodanda package in Python^70^ using a linear classifier based on a support vector machine architecture available within the scikit-learn package^71^. The following describes the fundamental steps implemented by Decodanda for the decoding analysis of neural activity. For details on the implementation, please refer to the package’s documentation: https://github.com/lposani/decodanda.

#### Data labeling

For all decoding analyses, we used inferred spiking activity as computed by CNMF. Each neuron’s spiking activity was z-scored based on the mean and standard deviation of its baseline activity from a 15-minute period prior to any CS presentation. Trials were obtained by taking time window of at least 500ms for tremble and stationary and windows of 2 seconds following ingress and were labeled according to the observed behavior and the presented stimulus (+1, +2, -1, -2). The average activity of each neuron within each trial was then computed to obtain a single vector of activity for each trial.

#### Pseudo-population sampling and cross-validation

The decoding performance was computed by a 20-fold cross-validation procedure. For each cross-validation fold, the trial activity vectors for each subject were randomly divided into training and testing sets (2/3 for training and 1/3 for testing) and used to construct two populations of training and testing pseudo-simultaneous (PS) activity vectors by pooling together all the recorded subjects that passed a minimum inclusion criterion of at least 4 trials per condition. To build a single PS activity vector, each subject was sampled *q* times to obtain vectors of length *qN*, where *N* is the total number of recorded neurons. In this work, we used *q* = 5. These pseudo-population sets were then used to train and test a linear support vector machine (SVM) to obtain a decoding accuracy value. The whole procedure, from training-testing separation to PS sampling and SVM accuracy testing, was repeated k=20 times, and the mean SVM decoding accuracy was taken as the measure of decoding performance.

#### Balanced sampling

Throughout the decoding analysis, the same number of PS vectors was sampled for all conditions (i.e. for all combinations of values of the decoded variables). Consider the decoding of two variables from the same neural data: *valence* (CS+ vs. CS-) and *behavior* (tremble vs. stationary). Since, in general, tremble is more frequent during CS+ than during CS-presentations, *valence* could act as a confound when decoding *behavior*, and vice-versa. This balanced sampling ensures that the four conditions (tremble & CS+, tremble & CS-, stationary & CS+, stationary & CS-) are equally represented in the training set. Thus, variables are ‘disentangled’, and their individual contribution to neural activity is analyzed independently of each other. Unless differently noted, throughout this work, we sampled 200 PS vectors for each condition. For comparing tremble and ingress decoding, in order to eliminate the possibility that activity related to a tremble extended beyond the tremble itself and thus affect the decoding of both tremble and ingress states, we restricted the analysis such that any trial could only contribute data for the decoding of ingress or tremble but not both. This restriction was achieved by randomly assigning all trials containing both trembles and ingress to the decoding of only one of the two response states.

### Decoding with a subsampled population of neurons

To analyze how decoding performance depends on the number of neurons used in the analysis, we repeated the decoding analysis, as described above, using a random subsample of the neural population with a given size n. We varied n in increments of 5% of the total number of neurons for a total of 19 values (5 to 100%). We repeated the random selection 10 times to obtain a population of decoding performances for each value of n.

### Cross-variable interference

We devised cross-variable interference (CVI) to measure the degree to which information about one variable (e.g. valence) can be used to decode a second variable (e.g. tremble) and vice-versa within a cross-validated decoding scheme. To compute the CVI, we used the same balanced cross-validated decoding scheme described above, with the difference that, across each cross-validation step, the training and testing data were labeled differently, each according to one of the two variables considered in the cross-variable analysis. The final CVI value was computed as the mean decoding performance across the two ways of choosing the training and testing variables (for example, training on tremble vs. stationary and testing on CS+ vs. CS-, and vice-versa). Note that each of these decoding performances is itself the average over k cross-validation folds.

### Cross-condition generalization performance

Cross-condition generalization performance (CCGP) was computed as described in Bernardi et al., 2020^27^. We first constructed pseudo-simultaneous activity vectors as described above, except we did not group data from pairs of conditions with the same decoding variable. Rather, trials used for training a given classification came from one of the pairs of conditions that both contained the decoding dichotomy for a given classification while sharing the same non-decoding variable. The corresponding testing set consisted of data from the other pair of conditions that shared the non-decoding variable. For example, when decoding *valence* across *behavior*, one training set consisted of data during interactions with CS+ versus CS-during *behavior*=tremble, and the testing set consisted of data from CS+ versus CS-during *behavior*=stationary. The decoding for a given dichotomy was then repeated, swapping the classes of trials used for the training and testing data. CCGP was obtained from the mean decoding performance from the two pairs of training and testing conditions.

### Null model and p-value

We tested the decoding performance and CCGP obtained by the procedures described above against a null model where the labels of trials were randomly shuffled. After each shuffling, the same procedures were repeated, obtaining a null-model value for decoding performance. We repeated the shuffling 10 times to obtain a distribution of null model performance values. The p-value for decoding performance or CCGP was then derived from the z-score of the performance computed on data compared to the distribution of null-model values.

### Multi-selectivity analysis

We performed the following analysis to assess whether cells were specialized for one of two variables (hereby called variable *A* and variable *B*) or whether they encode the variables with mixed selectivity. For a given variable *X* and each cell *i* we identified the coding importance of each individual cell, defined as 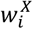, as the absolute value of its average decoding weight normalized by the standard deviation over *k* cross-validation folds:

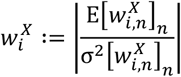

Where 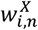 is the SVM decoding weight of cell *i* in the *n*-th cross-validation fold. We obtained a vector of all the values across the recorded population of cells:

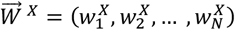

We denoted these vectors as 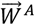, for variable *A* and 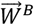 for variable *B*. If the recorded population is specialized, neurons that encode *A* will not encode *B*, and vice-versa. Therefore, a population of specialized neurons will be characterized by anti-correlated values of 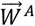 and 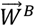 (Methods Figure 1a). On the other hand, if neurons are not specialized (mixed selectivity), we expect no relationship between *A* and *B* coding, resulting in a null correlation between 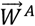 and 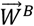 (Methods Figure 1b). A third possibility is that neurons are not specialized, but information is unevenly distributed across the population. In this case, neurons will typically encode neither or both variables, resulting in a positive correlation between 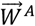 and 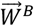 (Methods Figure 1c).

To assess whether the recorded population was specialized, we computed the Spearman correlation between the two coding importance vectors, denoted as 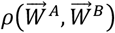 We then compared the value of 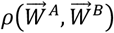 with those obtained by a null model where mixed selectivity is implemented by performing a solid random rotation of the coding weights of the two variables in the neural activity space. In this null model, the two coding vectors 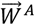 and 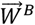 have no relationship with each other. For each null model iteration *k* we sample a random rotation matrix *R*_*k*_ and use it to rotate the weights vectors of A and B decoding before taking the average across cross-validations. This procedure was repeated 100 times, each time with a new rotation matrix, to obtain a population of null values. The recorded population of cells was then classified as either mixed or selective depending on the significance of the correlation, computed using the z-score of the Spearman correlation compared to the null model:

- 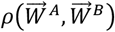< null model: selective population (Methods Figure 1a)
- 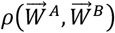∼ null model: mixed selectivity (Methods Figure 1b)
- 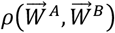> null model: mixed selectivity (uneven distribution of coding importance, Methods Figure 1c)

### L-score

In order to quantify the degree of specialization of a population of neurons for two given variables (hereby called variable A and variable B), we devised a measure of how much neurons lie on the axis in the weight-weight selectivity plot, which we called “L-score”. This measure is computed by measuring the overlap between the top *p* values for variable A and the top *p* values for variable B, normalized so that if the two sets are the same neurons the overlap is 1:

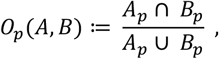

where *A*_*p*_ is the set of the top *p* selective neurons for variable A and *B*_*p*_ is the set of the top *p* selective neurons for variable B. By varying p from p=0 to p=1 (where *p* is expressed as fraction of the total number of neurons), we obtain a curve from which we compute the area under the curve, AUC(*O*_*p*_). If the selectivities for A and for B are uncorrelated random variables, the expected value of the overlap is 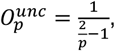, and the corresponding area under the curve is:

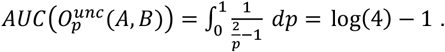

We then take the difference between the random uncorrelated case and the one computed from the data, 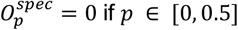, as a measure of specialization in the neural population. To obtain a value that ranges between 0 and 1 we then normalize with the expected value of AUC in the case of a perfectly specialized population, i.e., when half of the neurons have high selectivity for variable A and low selectivity for variable B and the vice-versa for the other half. In this specialized case 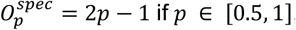 if *P* ∈ [0, 0.5] and 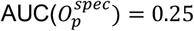 if *P* ∈ [0.5, 1], so the corresponding area under the curve is 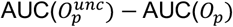 Finally, the L-score is defined as:

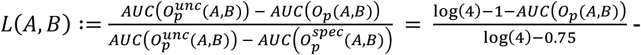

### Ablation analysis to assess decoding importance of selective cells

To assess the relative importance of specialized and mixed neurons in the decoding performance for a given variable, we performed an “ablation” analysis where each class of cells (specialized and mixed) is selectively excluded from the data used for the decoding analysis. Say we are testing the decoding importance of two variables, *A* and *B*. First, we identified the *n*_*B*_ specialized cells for variable *A* and the *n*_*A*_ specialized cells for variable *B* as described in the multi-selectivity analysis methods. Those neurons that are not specialized for either A or B were labeled as "mixed." We then excluded all the *n*_*A*_ + *n*_*B*_neurons that were identified as specialized and performed the decoding analysis for variables *A* and *B* (as described in the decoding methods) to get the decoding performances 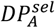 and 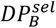 We then randomly selected the same number of mixed neurons by choosing a random value of selectivity lower than the selectivity threshold used to identify specialized neurons, which is equivalent to a random angle in the *W*_*A*_ vs. *W*_*B*_ graph (see Fig 3e and multi-selectivity methods), and choosing the *n*_*A*_ + *n*_*B*_ neurons that have the selectivity closest to this random value (pink dots in Fig. 3e). We excluded these neurons and repeated the decoding analysis to obtain the decoding performances 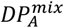 and 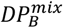. By repeating this random choice procedure 20 times, we obtained a population of decoding values for both variables when excluding mixed selectivity neurons from the decoding analysis (pink error bars in Fig. 3i). Finally, the decoding performances 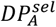 and 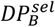 were compared to the population of 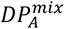 and 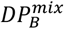 values to assess whether excluding specialized neurons affected decoding performance differently than excluding mixed-selective neurons. Statistical significance for each variable was assessed by computing the z-score of *DP*^*sel*^ compared to the corresponding *DP*^*mix*^ population.

### Visualization of neural patterns in a reduced-dimensionality space

To visualize the geometry of conditions in the neural activity space, we performed a dimensionality reduction of the recorded neural patterns using a PCA approach. Say we have *C* conditions. These could be, for example, the 2^ν^ combinations of ν experimental binary variables or multiple values of a single variable. First, for each condition, we sampled *N*_*c*_ pseudo-population patterns from the activity of the recorded subjects as described in the decoding analysis methods. We then used PCA to obtain the first 3 principal components of the resulting *C* ⋅ *N*_*c*_ sampled patterns. We then projected the *C* ⋅ *N*_*c*_ patterns on the 3 principal components to obtain *C* collections of *N*_*c*_ vectors in the three-dimensional PCA space. We then computed the centroid of each condition by taking the median position of its *N*_*c*_ vectors in the PCA space. Using the median instead of the mean ensures that outliers that could be due to artifacts will not dominate the position of the centroid. Finally, we visualized the resulting *C* centroids in the three-dimensional PCA space.

### Statistical Analysis

When paired comparisons were possible, paired t-tests were used to compute statistical significance. For comparing distributions, the Kolmogorov-Smirnov test was used to compute statistical significance. The alpha value for computing significance was 0.05. For all boxplots, the top and bottom edges of the box indicate the 75th and 25th percentile of the data. The whiskers extend to the most extreme data points not considered outliers. For decoder performance graphs showing shaded errors, the 95% confidence intervals obtained from the cross-validation runs are used for the upper and lower extents of the shading. Statistical analyses were performed using MATLAB (MathWorks) and Python.

---

**Supplemental Figure 1.**
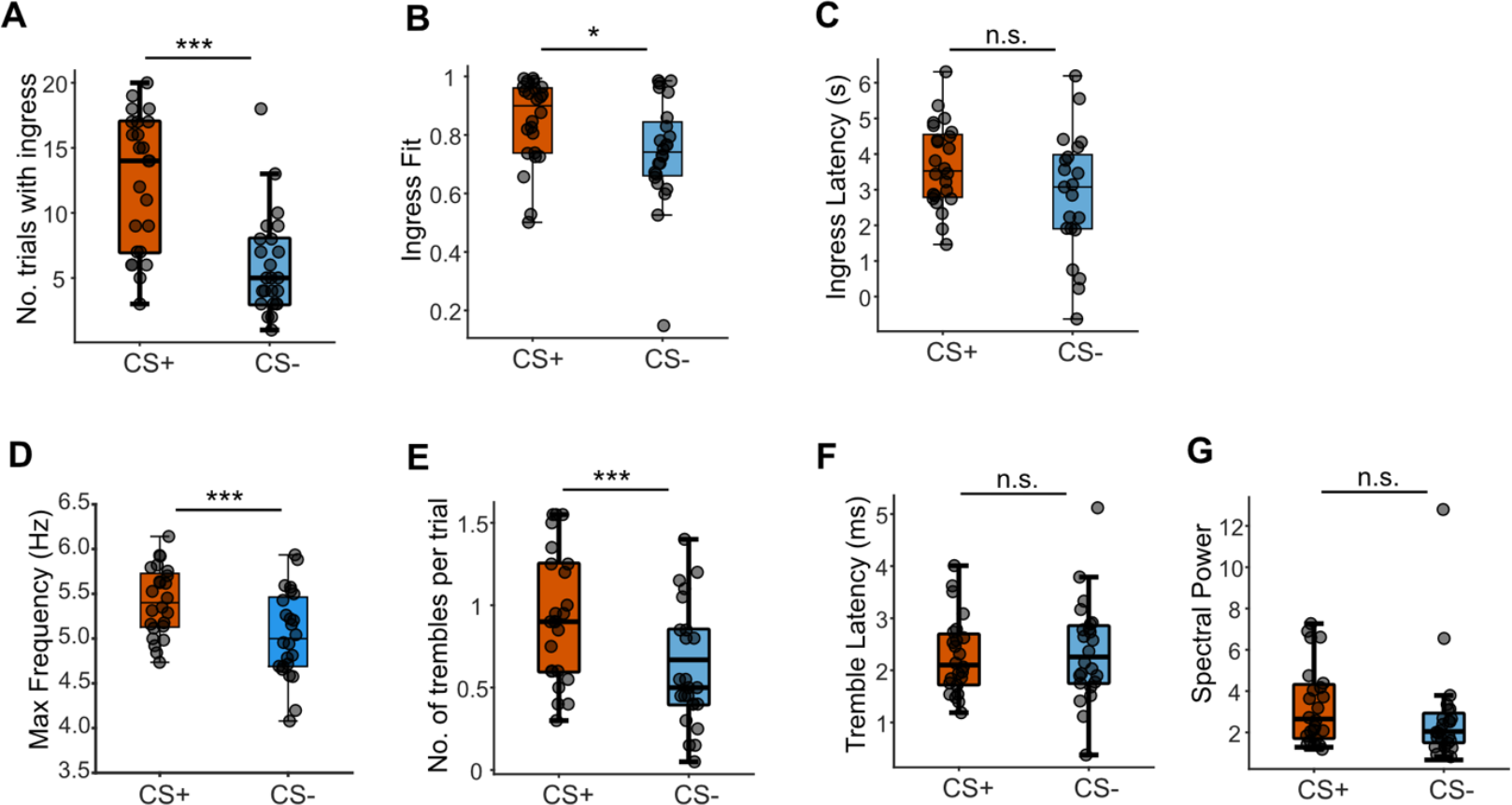
Quantification of CS+/- behavior in virtual burrow assay. **A**, Number of trials with ingress for CS+ and CS-trials a(cross mice (CS+, 12/20 trials; CS-, 6/20 trials) **B**, Ingress goodness of fit across mice (CS+ r2: 0.85, CS-r2: 0.74). **C**, Latency to ingress across mice (CS+: 3.5s CS-: 3.1s). **D**, Maximum frequency of a tremble bout across mice (CS+: 5.4Hz, CS-: 5.0Hz) **E**, Average number of trembles per trial across mice (CS+: 0.93, CS-: 0.61). **F**, Average latency to first tremble across mice (CS+: 2.3s CS-: 2.4s). **G**, Spectral power of tremble across mice (CS+: 3.3, CS-: 2.7). For all plots, n = 24 mice, *p<0.05, ***p<0.001, n.s. not significant, paired t-test.

**Supplemental Figure 2.**
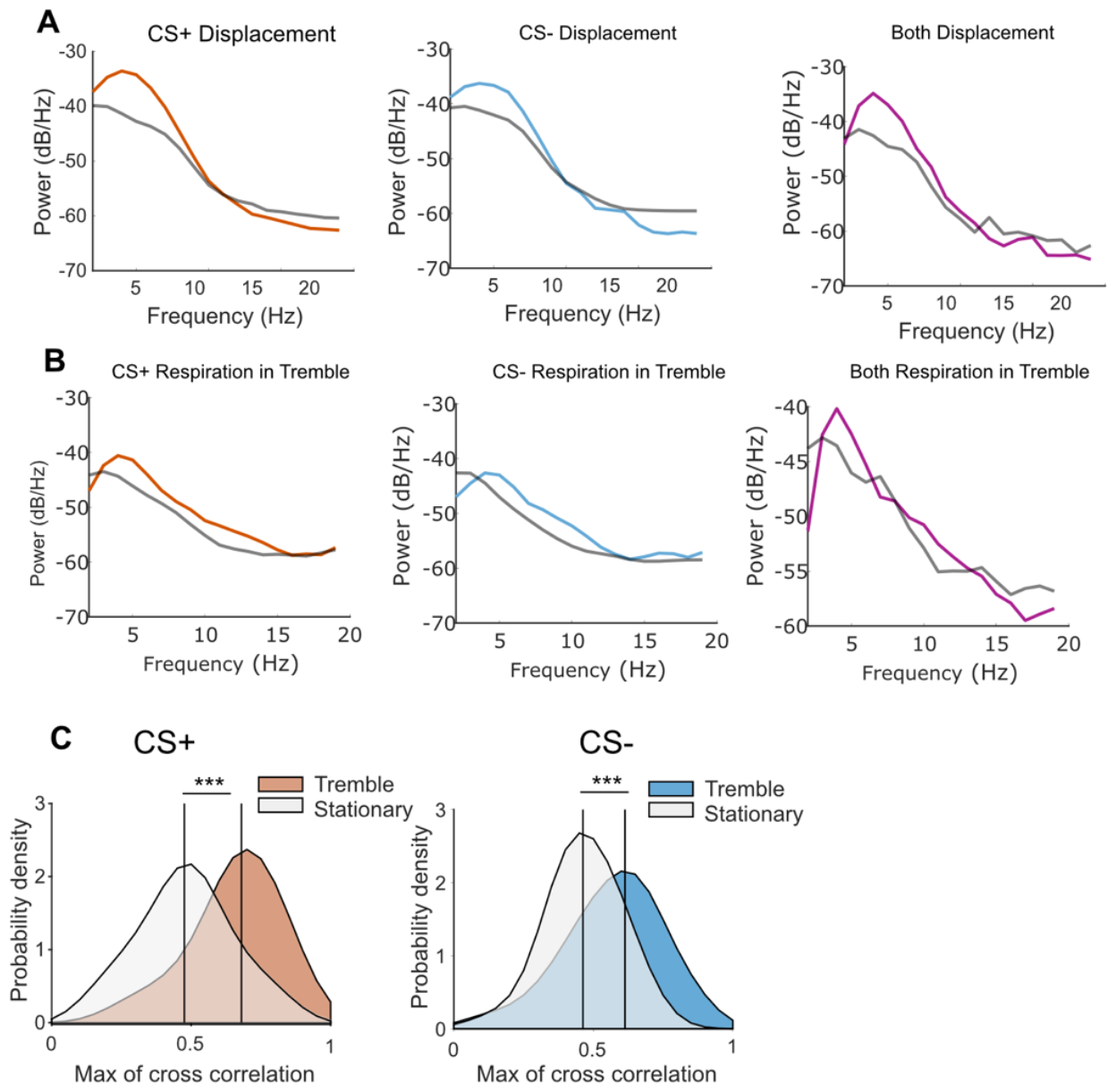
Quantification of the relationship between respiration and tremble. **A**, Average power spectra of tremble bouts in CS+ trials (orange), CS-trials (blue), and CS+/CS-trials combined (magenta) as compared to stationary bouts (gray). **B**, Average power spectra of respiration within tremble bouts in CS+ trials, CS-trials, and CS+ and CS-trials combined. **C**, Probability density distributions of the maximum correlation between displacement and respiration during tremble from CS+ (orange distribution) and CS-trials (blue distribution) vs stationary (gray distributions) separately (vertical lines depict median of distributions, ***p<0.001, Kolmogorov–Smirnov test).

**Supplemental Figure 3.**
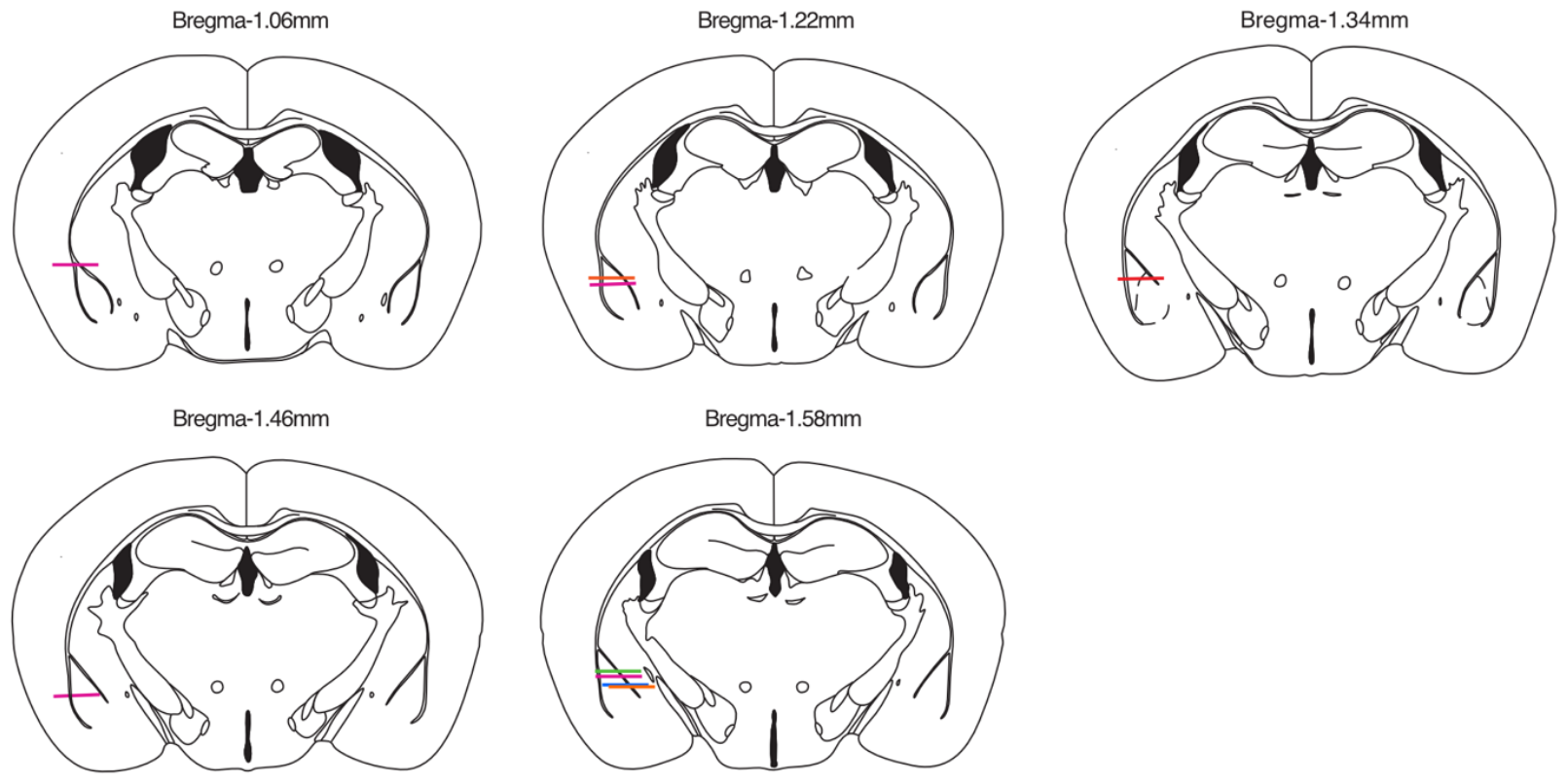
GRIN lens locations for mice in two photon imaging cohort. Each horizontal line depicts location of the GRIN lens implant within BLA of mice (n=9 mice) from subsequent histological examination.

**Supplemental Figure 4.**
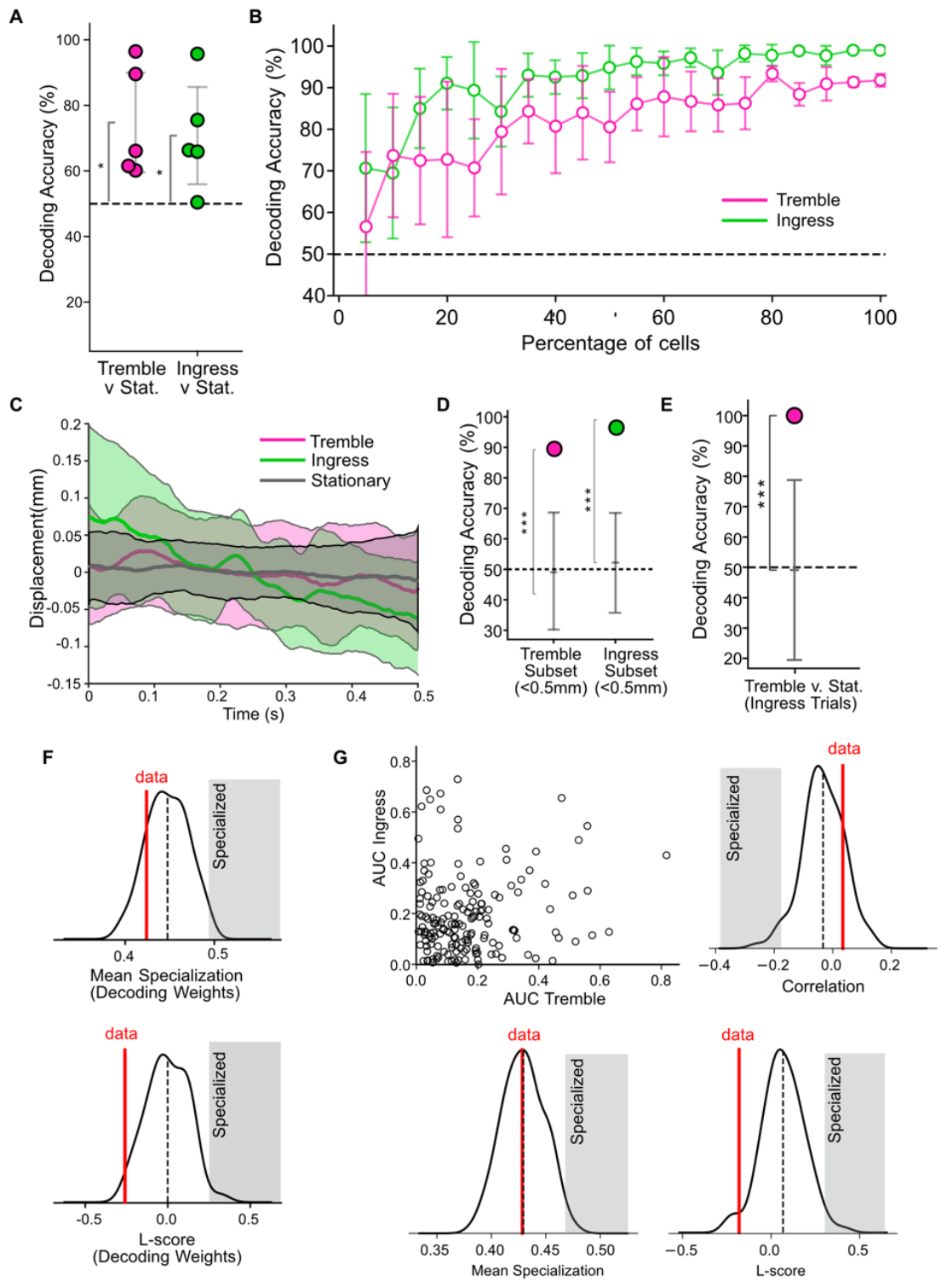
Individual animal decoding and number of neurons needed for decoding valence and identity. **A**, Decoding accuracy for tremble v. stationary and ingress v. stationary for each mouse (mean tremble: 75% accuracy, mean ingress: 71% accuracy, n=9 mice, *p<0.05, t-test from 50% chance accuracy). **B**, Decoding accuracy of tremble and ingress as a function of the percentage of neurons included in decoding analyses (resulting from analysis that down-samples neurons from the population). **C**, Subsets of tremble and ingress trials of at least 500ms with peak-to-peak displacement range less than 0.5 mm, similar to the displacement range of stationary epochs. Thick lines are mean across behavior category, shaded region depict ±sem. **D**, Decoding accuracy for tremble v. stationary and ingress v. stationary using only the subset of tremble or ingress trials with peak-to-peak displacement less than 0.5mm (tremble accuracy: 90%, ingress accuracy: 94%, ***p<0.001) **E**, Decoding accuracy for tremble v. stationary using only tremble bouts selected from trials with ingress (accuracy: 100%, ***p<0.001). **F**, Mean specialization and L-score of neurons calculated from normalized decoding weights from the data in Figure 3(E) (red line, ‘data’) and null model (distribution). **G**, For each neuron, the area under the curve (AUC) computed from an ROC analysis is plotted for both tremble and ingress. Spearman’s correlation, mean specialization, and L-score calculated between the AUC values for tremble and ingress data (red line, ‘data’) and from iterations of the null model (distribution).

**Supplemental Figure 5.**
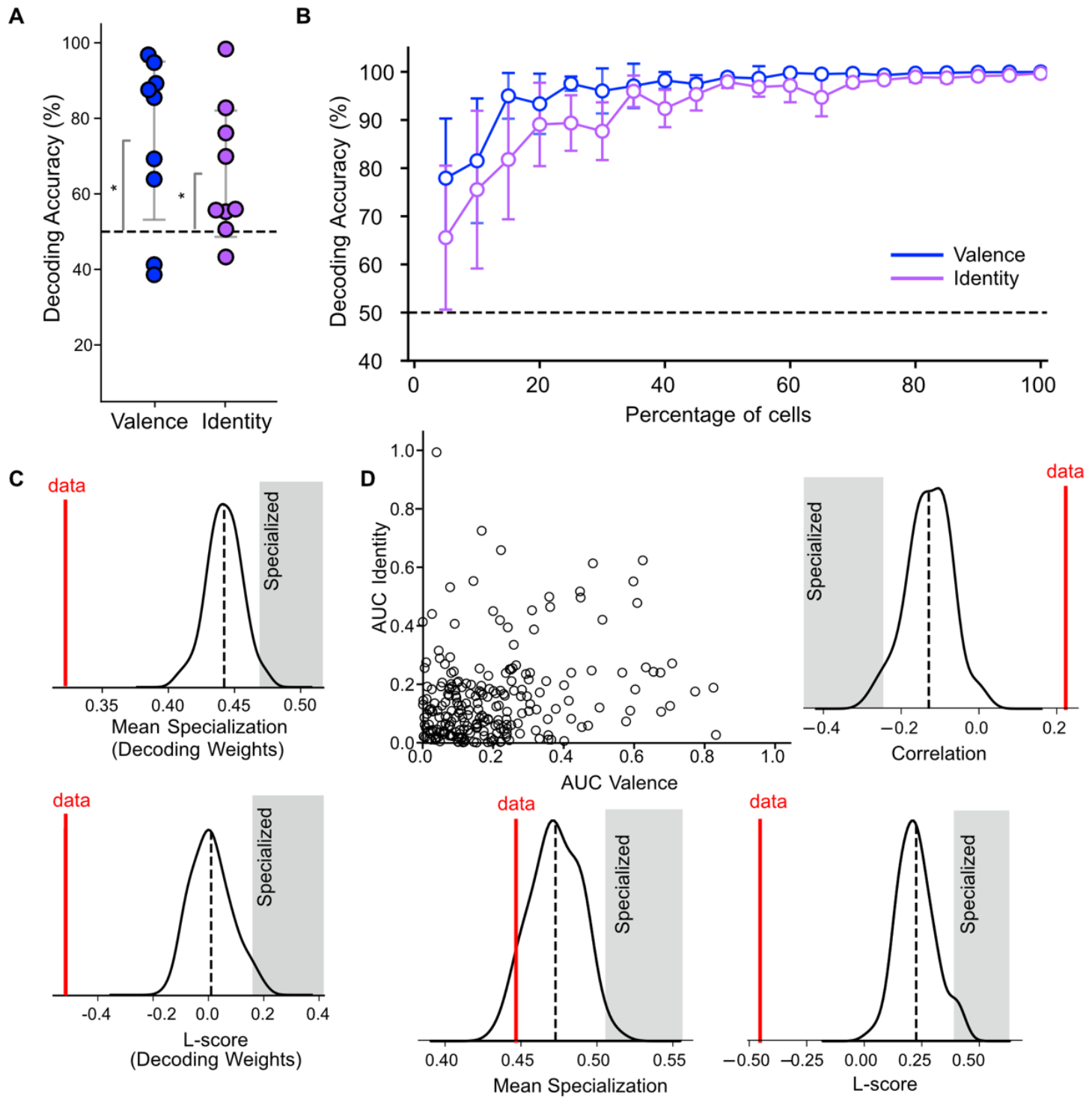
Individual animal decoding and number of neurons needed for decoding valence and identity. **A**, Decoding accuracy for CS valence and identity for each mouse (dots). Decoding accuracy for one of the two dichotomies related to CS identity (mean valence: 74% accuracy, mean identity: mean: 65% accuracy, n=9 mice, *p<0.05, t-test from 50% chance accuracy) **B**, Decoding accuracy of valence and one of the two identity-related dichotomies (CS+1/CS-1 vs CS+2/CS-2) as a function of the percentage of neurons included in decoding analyses (resulting from analysis that down-samples neurons from the population). **C**, Mean specialization and L-score of neurons calculated from normalized decoding weights from the data in Figure 4(D) (red line, ‘data’) and null model (distribution) **D**, For each neuron, the area under the curve (AUC) computed from an ROC analysis is plotted for both valence and identity. Spearman’s correlation, mean specialization, and L-score calculated between the AUC values for valence and identity data (red line, ‘data’) and from iterations of the null model (distribution).

**Supplemental Figure 6.**
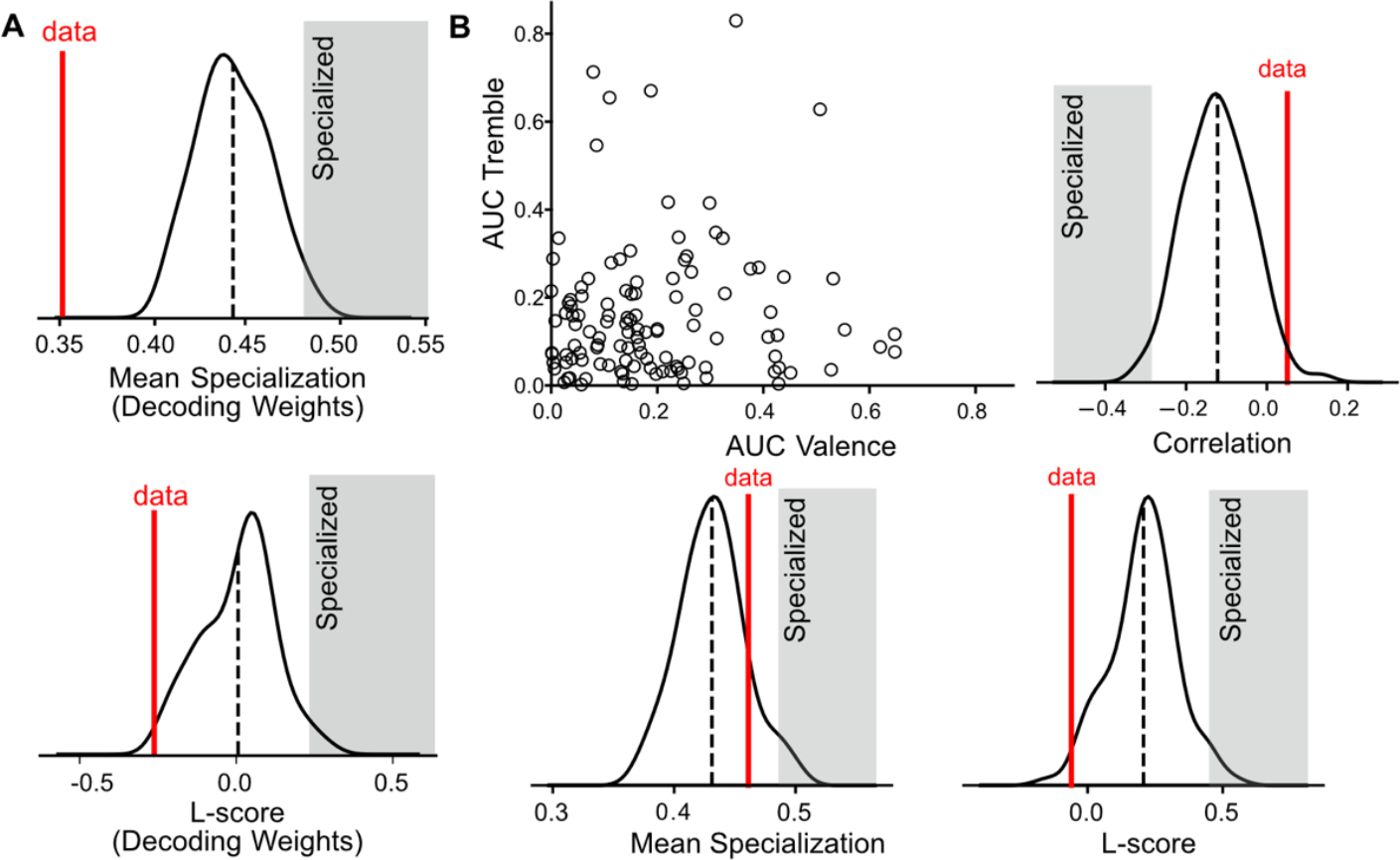
BLA mixed selectivity for valence and tremble characterized using decoding weights and ROC analysis. **A**, Mean specialization and L-score of neurons calculated from normalized decoding weights from the data in Figure 5(C) (red line, ‘data’) and null model (distribution) **B**, For each neuron, the area under the curve (AUC) computed from an ROC analysis is plotted for both valence and tremble. Spearman’s correlation, mean specialization, and L-score calculated between the AUC values for valence and tremble data (red line, ‘data’) and from iterations of the null model (distribution).

**Supplemental Figure 7.**
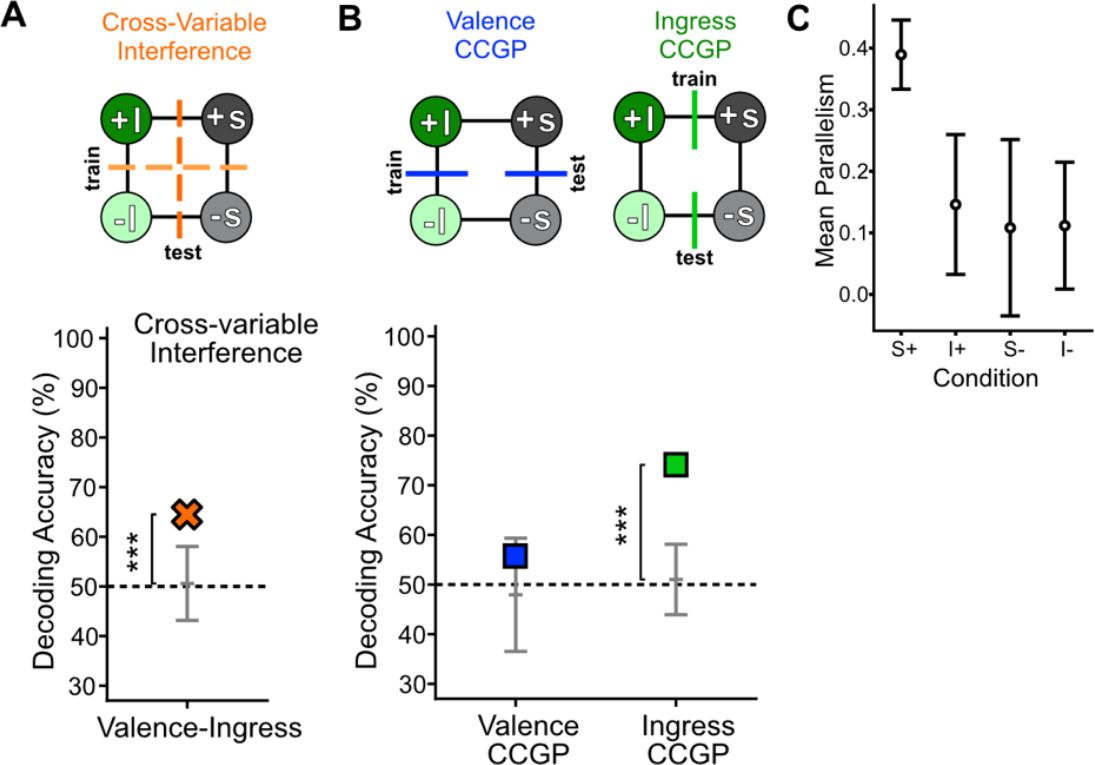
CCGP and cross-variable interference for ingress and valence. **A**, Decoding performance for Cross-Variable Interference between ingress and valence is above chance (accuracy: 64%, ***p<0.001), indicating that the coding directions for these variables are not orthogonal. **B**, BLA neural ensembles exhibit a geometry where CCGP for valence is chance but CCGP for ingress is high (valence CCGP: 56% accuracy, ingress CCGP: 74%, ***p<0.001; grey error bars, 95% confidence intervals for the null model). **C**, Mean parallelism score computed between coding directions between each condition and each of the other three conditions (see Methods). The coding direction between the +S (unsafe) condition and each of the other (safe) conditions have a high parallelism score, accounting for high CCGP in Fig. 7F. This type of geometry (parallel coding directions) is not observed with respect to the other conditions (-S, - I, +I).

